# Emergent Actin Flows Explain Diverse Parasite Gliding Modes

**DOI:** 10.1101/2022.06.08.495399

**Authors:** Christina L. Hueschen, Li-av Segev Zarko, Jian-Hua Chen, Mark A. LeGros, Carolyn A. Larabell, John C. Boothroyd, Rob Phillips, Alexander R. Dunn

## Abstract

During host infection, single-celled apicomplexan parasites like *Plasmodium* and *Toxoplasma* use a motility mechanism called gliding, which differs fundamentally from other known mechanisms of eukaryotic cell motility. Gliding is thought to be powered by a thin layer of flowing filamentous (F)-actin^1–3^ sandwiched between the plasma membrane and a myosin-coated^4,5^ inner membrane complex. How this surface actin layer drives the diverse apicomplexan gliding modes observed experimentally - helical, circular, and twirling^6,7^, and patch^8^, pendulum^9^, or rolling^2^ – presents a rich biophysical puzzle. Here, we use single-molecule imaging to track individual actin filaments and myosin complexes in live *Toxoplasma gondii*. Based on these data, we hypothesize that F-actin flows arise by self-organization, rather than following a microtubule-based template as previously believed. We develop a continuum model of emergent F-actin flow within the unusual confines provided by parasite geometry. In the presence of F-actin turnover, our model predicts the emergence of a steady-state mode in which actin transport is largely rearward. Removing actin turnover leads to actin patches that recirculate up and down the cell, a “cyclosis” that we observe experimentally for drug-stabilized actin bundles in live parasites. These findings provide a mechanism by which actin turnover governs a transition between distinct self-organized F-actin states, whose properties can account for the diverse gliding modes known to occur. More broadly, we illustrate how different forms of gliding motility can emerge as an intrinsic consequence of the self-organizing properties of F-actin flow in a confined geometry.

---

Single-celled parasites of the eukaryotic phylum Apicomplexa cause hundreds of millions of cases of malaria, toxoplasmosis, and cryptosporidiosis each year^10–12^. To propel themselves over host cells and through extracellular matrix, motile Apicomplexa like *Plasmodium* spp. or *Toxoplasma gondii* use an adhesion-dependent locomotion mechanism called gliding that defies the paradigmatic classification of eukaryotic cells into cilia-dependent swimmers and cell-shape-change-dependent crawlers. Gliding depends on a layer of F-actin^1,2^ and a fast, single-headed myosin, MyoA^4^, confined to a 25 nm-thick compartment between the parasite plasma membrane and a membranous scaffold termed the inner membrane complex (IMC)^13^. MyoA is anchored in the inner membrane complex through its association with myosin light chain 1 (MLC1)^5,13^. In one prevalent mechanistic model, MyoA slides short actin filaments rearward through the inter-membrane space, toward the posterior end of the cell (reviewed in ^3,14^). When actin-coupled adhesin proteins in the plasma membrane bind to a stationary external substrate, MyoA instead propels the inner cytoskeleton and parasite cytoplasm forward. In this model, the rearward direction of actin filament transport by MyoA is thought to be fixed, and likely templated by a basket of polarized subpellicular microtubules beneath the inner membrane complex^3^. However, as discussed below, this “templated” model cannot account for all observed apicomplexan gliding motions.

On a two-dimensional substrate, motile *Toxoplasma gondii* parasites can undergo helical gliding, with simultaneous forward translation and cell body rotation^6^ (**Video 1**), or glide in circles with their anterior end leading, motions that translate into a corkscrew trajectory when embedded in three-dimensional matrix^15–17^. In addition, parasites display a rotational motion known as twirling when oriented upright, with their posterior end on the substrate. These unidirectional cell movements contributed to the working model of rearward actin transport along the path of subpellicular microtubules^3^. However, observations of back-and-forth motion, termed patch or pendulum gliding, have been reported in a diversity of conditions (**Table S1, Fig. S1A, Video 1**), and prevalent models cannot explain these motions. Further, MLC1-MyoA complexes localize throughout the inner membrane complex, not merely above subpellicular microtubules^18,19^ (**Fig. S1B**), making it unclear how myosin orientation might be fixed relative to the cell axis. The recent discovery of gliding by *Plasmodium* merozoites^20^, which do not have a basket of chiral subpellicular microtubules, further suggests that our understanding of how gliding motility arises from molecular-level organization remains incomplete. In this study, we sought to better understand how the actomyosin machinery gives rise to such a diverse array of gliding movements, and more broadly, how actin and myosin are patterned or polarized to yield coherent force generation in this system.

## Dynamics of single actin and myosin proteins in motile *Toxoplasma gondii*

To directly test the model of “templated” rearward actin transport along the path of subpellicular microtubules, we used single-molecule fluorescence imaging^21,22^ to track individual labeled actin (ACT1) and myosin light chain 1 (MLC1) proteins in live, active *Toxoplasma gondii* tachyzoites (**Fig. 1A**). Pauses between parasite movements enabled us to reliably track individual molecules (see Methods) relative to microtubule polarity, which is fixed through the tachyzoite life stage. MLC1 molecules were frequently immobile (“bound”) for tens of seconds (**Fig. 1B, Fig. S2A, Video 2**), consistent with a stably-anchored population of MyoA motors at the inner membrane complex. Relative to MLC1, a larger fraction of actin molecules were mobile (**Fig. S2B**) and displayed meandering (“diffusive”) behavior as well as ballistic (“directional”) behavior consistent with processive transport by myosin (**Fig. 1C, Video 3**). Directional actin molecules moved with a mean speed of 4.8 µm s^-1^ (**Fig. 1D**), similar to the *in vitro* actin transport speeds of 4-5 µm s^-1^ reported for purified *Toxoplasma* MyoA complexes^23,24^.

**Figure 1.**
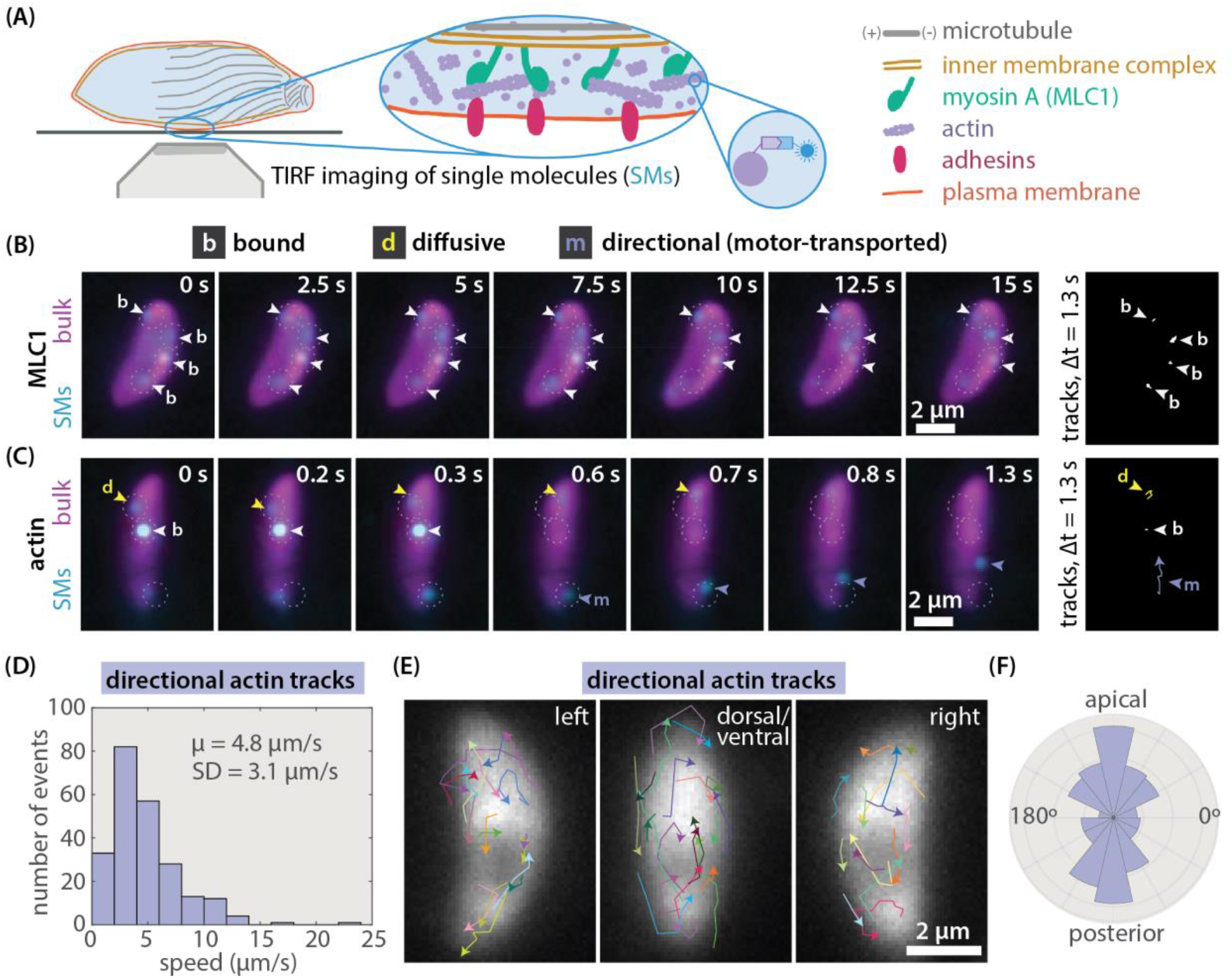
*Toxoplasma gondii* actin transport direction is heterogeneous, not uniformly rearward. (**A**) Schematic of the inter-membrane actomyosin layer that drives apicomplexan gliding and of total internal reflection fluorescence (TIRF) imaging of single molecules (SMs) of halo-actin or MLC1-halo (low expression; sparse labeling with Janelia Fluor dyes (cyan)). (**B**)-(**C**) Examples of myosin (MLC1) and actin single molecule dynamics (cyan) over time in extracellular parasites, with bulk labeling (magenta) to show cell position. Arrowheads highlight examples of specific molecule behaviors. Single molecule trajectories (far right) are shown for an equal time interval (1.3 s) to allow comparison of bound (white), diffusive (yellow), and directional (purple) movements. (**D**) Histogram of speeds of directional actin tracks from 18 cells. (**E**) Directional actin tracks from 18 overlaid cells from 3 experiments, aligned with anterior end up and grouped by the cell side visible. Cell polarity was determined by microtubule labeling or tracking of the posterior end following posterior-down cell twirling (Supplemental Information), and directional actin tracks were aligned with respect to the parasite long axis. **(F)** Polar histogram of the orientation of directional actin displacements with respect to cell polarity (n = 231 displacements, 54 tracks, 18 cells).

Analysis of directional actin tracks in live parasites was inconsistent with a mechanistic model featuring fixed myosin polarity and uniformly rearward actin flow. Relative to the parasite long axis, F-actin transport direction was heterogeneous and as often forward as rearward (**Fig. 1E, F)**. While subpellicular microtubule orientation could explain the observed bias toward longitudinal F-actin transport (**Fig. 1F**), for example by templating closer longitudinal than latitudinal MyoA spacing^25^, microtubule polarity evidently did not dictate rearward-only actin transport.

## A theoretical model of actin filament collective motion predicts emergent actin organization

Our data, combined with the diversity of gliding modes exhibited by motile Apicomplexa, led us to explore the possibility that gliding motility might represent an emergent self-organized state rather than the consequence of a microtubule-templated asymmetry in actomyosin polarity. In this scenario, the heterogenous F-actin transport observed between glides (**Fig. 1D-F**) could reflect a disorganized state, between transient self-organized actin states that drive gliding. Self-organization^26^ is a hallmark of actomyosin networks, with morphologically diverse examples such as the lamellipodia of crawling keratocytes^27^ and neutrophils^28^, the flowing cortex of *C. elegans* zygotes^29^, and dense actin networks of *in vitro* motility systems^30^. Drawing on continuum theories for active collective motion, or flocking^31–33^, and previous studies of self-organization of cytoskeletal systems^34–36^, we developed a continuum model of *Toxoplasma* actin filament collective motion (**Fig. 2; Fig. S3**). In our model, actin filaments at the surface of the cell follow a few simple rules: filaments are transported at the speed of myosin motors, and filament orientation sets the myosin-driven transport direction (**Fig. 2A**); filaments align with neighboring filaments through collisions^37^ or due to crosslinking proteins like *Toxoplasma* coronin^38–40^ (**Fig. 2B**); filament density remains within a realistic range (**Fig. 2C**); and filament alignment is biased away from orientations of high membrane curvature (**Fig. 2D**). Actin filament organization is described by two fields: the scalar field *ρ*, which captures filament density, and the velocity vector field **v**, which captures both filament polarity (orientation of **v**) and speed (magnitude of **v**). Two equations are needed: the continuity equation

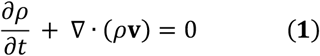

ensures conservation of filaments, and filament velocity evolves according to the rules described above using the minimal Toner-Tu equations

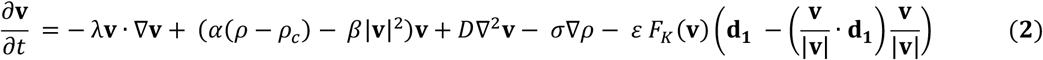

where *ρ*_c_ is the critical density above which filaments move coherently, the coefficient *λ* tunes filament transport (self-advection), the ratio of α and β sets a filament transport speed scale, D tunes filament alignment with neighbors, σ∇ρ provides an effective pressure that limits density variance, and the coefficient ε tunes the curvature-induced force F_κ_(**v**) that rotates filaments away from **d**_**1**_, the direction of maximum curvature (**Fig. S3**). The exclusion of any of these terms leads to results that are either aphysical or inconsistent with established *Toxoplasma* biology (Supplementary Information). While likely a simplification, the generality of this minimal flocking framework allowed us to explore how cell-scale actin organization might emerge from local actomyosin interactions confined to *Toxoplasma’*s surface shape.

**Figure 2.**
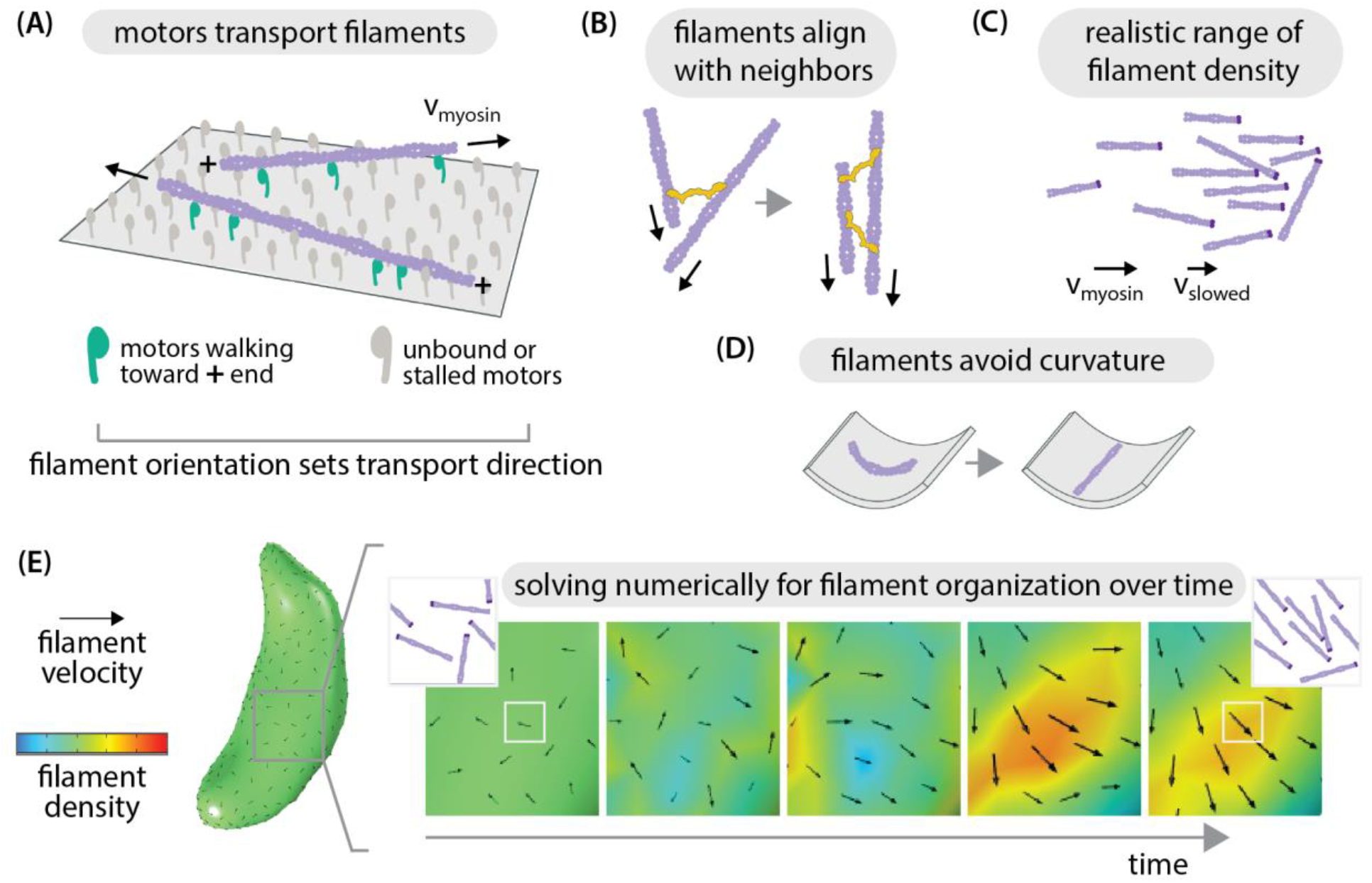
*Toxoplasma* actin self-organization: theoretical model. Rules of local actin filament behavior, implemented in equation (2): (**A**) Actin filaments (purple) are transported with their minus ends leading at speed *v*_*myosin*_ as indicated by arrows; Neighboring filaments align by steric effects or crosslinking proteins (yellow); (**C**) filament density remains within a realistic range, with filament speed slowing if entering a pile-up; (**D**) filament orientation is biased toward lower curvature. (**E**) Example of numerically solving for filament self-organization using the finite element method, predicting filament density and velocity over time. Black arrow size reflects velocity magnitude.

To predict what cell-scale actin organization patterns could emerge from the molecule-scale rules illustrated in **Figure 2**, we began with a disordered network and asked how filament density and velocity evolve over time (**Fig. 2E**), using the finite element method to solve equations (1) and (2) in COMSOL Multiphysics®. Importantly, we sought to incorporate the true shape of this thin membrane-constrained layer of actin, which can be approximated as a two-dimensional closed surface following the rigid and stereotypical shape of *Toxoplasma gondii* tachyzoites. We used soft X-ray tomography^41^ to obtain native-state high-resolution 3D reconstructions of cryo-fixed extracellular parasites and used a spherical harmonic description^42^ to convert them to closed surfaces for finite element method analysis (**Fig. 3A**). We then derived a curved-surface formulation of our governing equations (1) and (2) using extrinsic differential geometry^43–45^ (Supplementary Information). In essence, the resulting formulation uses the surface normal vector to project into the local tangent plane, and thus requires no intrinsic surface parameterization. We note the versatility of such an approach for solving continuum models on complex geometries for both living and non-living systems^44^.

**Figure 3.**
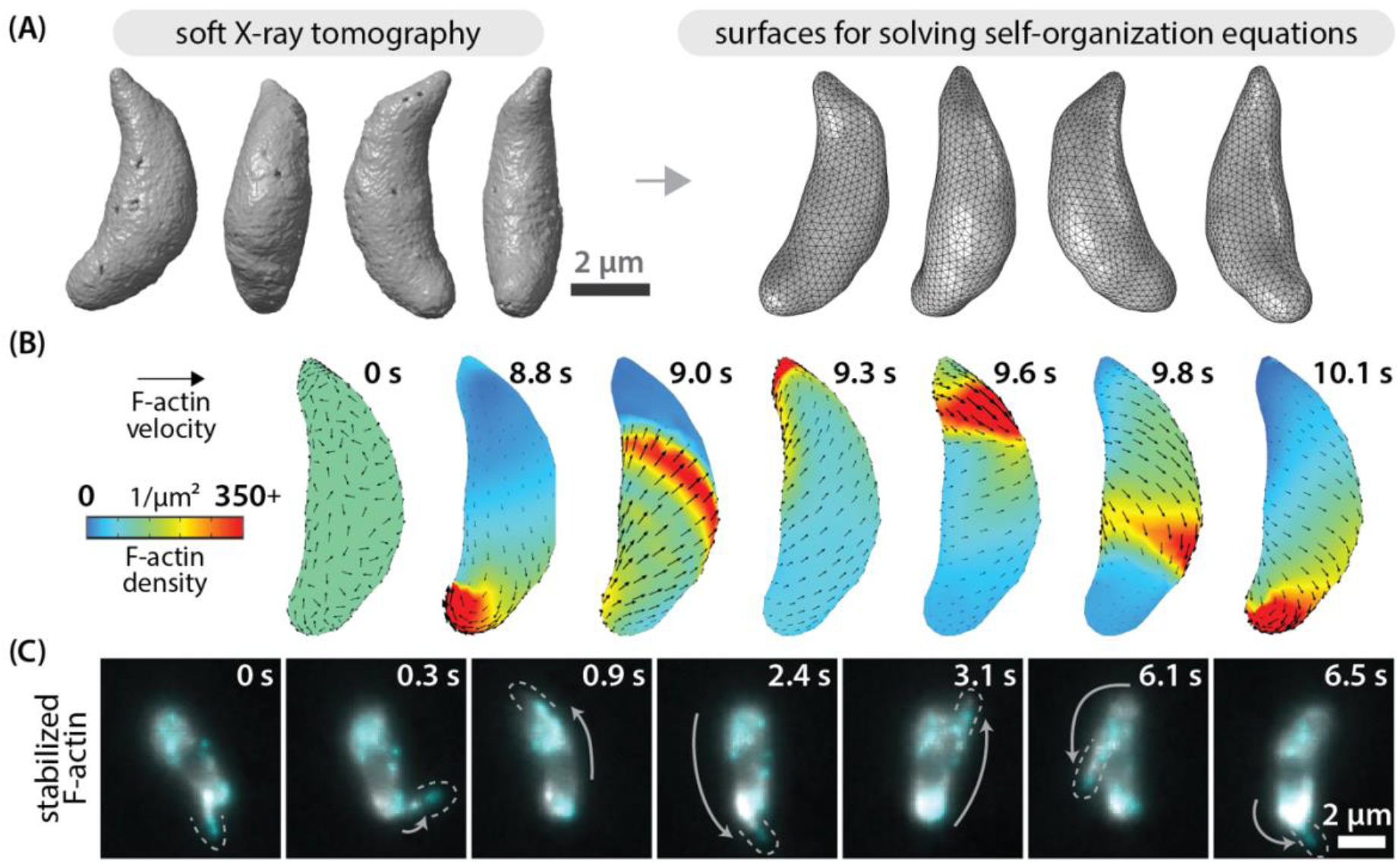
Stable actin filaments circle the *Toxoplasma* cell. (**A**) Soft x-ray tomograms of cryo-fixed extracellular *Toxoplasma gondii* tachyzoites were used to generate triangle-meshed surfaces on which to solve our actin self-organization theoretical model. (**B**) For stable filaments, solving the model constrained to *Toxoplasma*’s surface geometry predicts recirculating actin patches. (**C**) In experiments, jasplakinolide-stabilized actin filaments can circle around the cell. Cyan and gray both show actin, labeled at different dye densities. Dotted lines outline protruding actin filaments.

### Polarized actin turnover governs a transition between recirculation and unidirectional actin transport

Starting from a disordered initial state and, importantly, the assumption of a conserved number of stable actin filaments enforced by equation (1), numerical simulations predicted the emergence of patches of parallel actin filaments that circulate up and down along the cell as shown in **Figure 3B** and **Video 4**. We observed a similar recirculation of F-actin in experiments, imaging actin bundles in *Toxoplasma* tachyzoites treated briefly with the actin-stabilizing drug jasplakinolide (**Fig. 3C, Video 5**). Thus, in the absence of filament turnover, a self-organization model predicts the emergence of F-actin recirculation – a continuous “cyclosis” observed experimentally for stabilized filaments in live parasites.

Next, we extended these theoretical and experimental results to consider regimes of filament turnover. Importantly, *Toxoplasma* helical and circular gliding modes are known to require regulated actin depolymerization by proteins like profilin^46^ and actin depolymerizing factor (ADF)^47^. Further, the polymerization of F-actin essential to gliding depends on formin 1 (FRM1), which localizes to the parasite anterior^48,49^. Estimates of F-actin lifetime and the characteristic timescale of F-actin cyclosis are both on the order of seconds, justifying an addition of filament turnover to our theoretical model that extends it beyond prior flocking models^50^ (Supplementary Information). Based on current knowledge, polymerization is favored specifically at the cell anterior, while depolymerization by profilin and ADF is not known to be spatially restricted. Thus, in the filament turnover model, F-actin density at the anterior cell surface is governed by

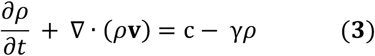

where c tunes F-actin polymerization and stabilization (rate of filaments produced per unit area), and *γ*tunes depolymerization (rate of filament loss). Outside the cell anterior (**Fig. S3**), F-actin density is governed by

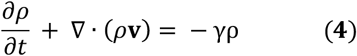

while equation (2) governs F-actin velocity across the entirety of the cell.

The addition of F-actin turnover and anterior polymerization enabled the emergence of a new F-actin organization state, in which actin transport is largely unidirectional and rearward (**Fig. 4A, Video 6**). In this emergent state, filament density and velocity reach a steady-state: while individual actin filaments flow continuously rearward, the average F-actin density and velocity at a given position reaches a fixed value. Tuning filament polymerization and depolymerization rates (**Fig. 4B)** shifted the emergent F-actin pattern between states. Increasing anterior polymerization and increasing filament stability (lowering depolymerization rate) favored the F-actin cyclosis described in **Figure 3**. Conversely, increasing filament depolymerization rate favored unidirectional F-actin transport. At an intuitive level, the transition from cyclosis to unidirectional flow occurs as filament lifetime (1/*γ*) drops below the cross-cell filament transport time (∼*L*_*cell*_/*v*_*myosin*_), preventing a posterior pile-up of F-actin large enough to force recirculation. In summary, actin turnover governs a transition between two self-organized states: F-actin recirculation and steady-state unidirectional transport.

**Figure 4.**
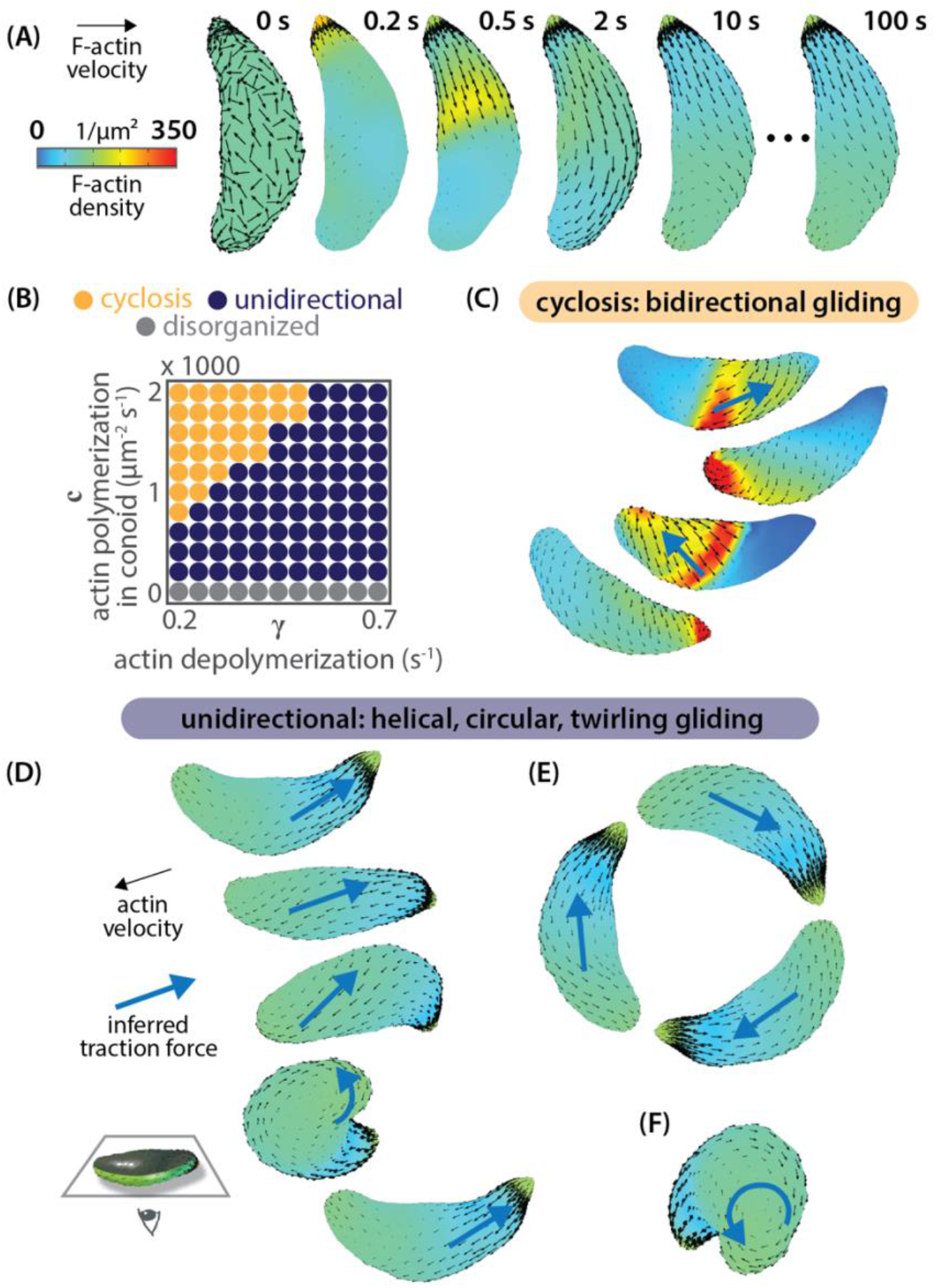
Polarized actin turnover governs a transition between actin recirculation and unidirectional transport. (**A**) Incorporating F-actin depolymerization and apical polymerization into the model enables the emergence of a unidirectional, stable velocity pattern. (**B**) Tuning rates of F-actin polymerization and depolymerization move the cell between distinct self-organized states. Model prediction: recirculating F-actin cyclosis generates bidirectional traction force (blue arrows) to drive “patch” gliding. (**D**) Model prediction: the unidirectional self-organized F-actin state drives helical gliding, (**E**) circular gliding, and (**F**) twirling. In the images in (C)-(F), cells are viewed from below; cell-substrate contact occurs at the position of the inferred traction force.

## Distinct self-organized actin states can account for diverse gliding behaviors

In this section, we develop the hypothesis that actin self-organization into distinct states explains the rich diversity of apicomplexan cell movements observed experimentally, from helical gliding and twirling to back-and-forth patch gliding^2,6,8^. During gliding on a surface, parasites form an adhesive cell-substrate contact point, where adhesin proteins bind the external substrate and form a stationary patch^49,51^. Cell motion occurs when MyoA walks the inner cytoskeleton and parasite cytoplasm toward the plus ends of F-actin adhering to that stationary patch^3,23^. Therefore, local F-actin polarity dictates the direction of myosin-powered traction force and the direction of cell movement. A map of self-organized actin velocity (**Fig. 3B, 4A**) thus implies a corresponding map of traction force direction (blue arrows, **Fig. 4C-E**).

In the recirculating actin state, a qualitative translation of predicted F-actin velocity patterns into traction force orientation (blue arrows, **Fig. 4C**) can explain the previously puzzling observations of back-and-forth *Toxoplasma* and *Plasmodium* cell gliding summarized in **Table S1**. These observations include patch gliding, pendulum gliding, and rolling in conditions like *Toxoplasma gondii* actin depolymerization factor (ADF) knockout cells^47^, *Toxoplasma gondii* treated with actin stabilizers^2^, and *Plasmodium berghei* sporozoites with mutations in the actin-binding domain of the adhesin protein TRAP^9^. Our theoretical finding that increased filament stability shifts F-actin self-organization from a unidirectional to recirculating mode (**Fig. 4B**) provides a unifying interpretation of these disparate experimental results (**Table S1**).

In the unidirectional regime, tuning the rate of F-actin depolymerization changes features of the predicted velocity patterns, including chirality and density gradient length scale (**Fig. S4**). For choices of polymerization and depolymerization rate close to the unidirectional-to-recirculating transition, emergent F-actin velocity patterns are consistent with the observed mechanics of helical gliding, circular gliding, and twirling (**Fig. 4D-F**). Helical gliding initiates when the ‘left’ side of the cell is in contact with the substrate, while circular gliding initiates given ‘right’ side contact (considering the concave cell surface to be ventral)^6^. In each case, inferred traction force vectors along the path of substrate contact can explain observed cell translation and rotation (**Fig. 4D**,**E**). Similarly, predicted vortical F-actin polarity at the parasite posterior would lead to a myosin-powered torque and cell rotation or twirling, which is indeed characteristic during cell posterior contact (**Fig. 4F**). Thus, we hypothesize that the cyclosis mode of F-actin self-organization (**Fig. 3B, 4C**) drives bidirectional cell gliding (patch, pendulum, rolling), the unidirectional mode of F-actin self-organization (**Fig. 4A,D-F**) drives helical gliding, circular gliding, and twirling, and that actin turnover governs the transition between modes.

### Outlook

We hope that the theoretical and experimental results presented here will prove a stimulating first step in understanding actomyosin self-organization in the Apicomplexa, given the fruitfulness of the self-organization paradigm as a null hypothesis for cytoskeletal systems across biology. Far from wishing to claim finality for the particulars of the model developed here, we look forward to the incorporation and discovery of additional biological complexity through a dialogue between theory and experiment. Such a dialogue will benefit from fast and sensitive volumetric imaging of actin single molecules, combined with sufficiently sophisticated analysis algorithms, to enable 3D reconstructions of F-actin velocity fields in the reference frame of the cell *during* specific gliding motions, and from quantitative comparison of predicted and measured traction forces and gliding mechanics^8,51^.

Broadly, we note continuum theory’s ability to unify natural phenomena across scales, allowing a flocking theory inspired by collective bird motion^31,33^ to provide insight into microscopic actin organization in a unicellular parasite. Looking forward, the mathematical framework developed here will enable a meaningful examination of actomyosin self-organization in *Plasmodium* spp. sporozoites, ookinetes, and other motile Apicomplexa, incorporating their characteristic cell shapes to generate self-organized patterns of actin velocity and inferred traction force for comparison to experimental data. Further, reports of gliding cells exist within at least three major clades of eukaryotic life^52,53^, suggesting that this “esoteric” mode of cell locomotion may in fact be common but understudied, and deserving of a unifying effort to understand its physical principles and their degree of conservation across Eukarya.

## Supplementary Information

Figures S1-S8 Videos 1-6

Supplementary Table 1

Experimental Methods and Materials

Theoretical Model

Supplementary References

## Supporting information

Video1-GlidingModes

Video2-MyosinSingleMolecules

Video3-ActinSingleMolecues

Video4-CyclosisPrediction

Video5-StableBundleCyclosis

Video6-UnidirectionalPrediction

## Acknowledgments

We are grateful for helpful discussions with colleagues and friends. We thank in particular Melanie Espiritu, Greg Huber, Elgin Korkmazhan, Madhav Mani, Mike Panas, Manu Prakash, Carlos Rojo, Suraj Shankar, Sho Takatori, Yuhai Tu, Vipul Vaccharajani, the Stanford Apicomplexa Supergroup, and members of the Dunn lab. Gary Ward, Rachel Stadler, and Deepak Krishnamurthy were invaluable sources of inspiration and guidance throughout. This work was supported by a Damon Runyon Fellowship Award (C.L.H.), a Burroughs Wellcome Career Award at the Scientific Interface (C.L.H.), NIH R35GM130332 (A.R.D.), an HHMI Faculty Scholar Award (A.R.D.), NIH MIRA 1R35 GM118043 (R.P.), and the Chan Zuckerberg Biohub Intercampus Team Award (J.C.B., C.A.L.). The soft X-ray tomography was conducted at the National Center for X-ray Tomography, which is supported by NIH NIGMS (P30GM138441) and the DOE’s Office of Biological and Environmental Research (DE-AC02-5CH11231). The Center is located at the Advanced Light Source, a U.S. DOE Office of Science User Facility under contract no. DE-AC02-05CH11231.

## Author Contributions

C.L.H., A.R.D., R.P., L.S.Z., and J.C.B. conceptualized the study; C.L.H., L.S.Z., and J.H.C. performed experiments; C.L.H. analyzed data with insight from A.R.D., R.P., and J.C.B.; C.L.H. and R.P. developed the theoretical model and performed computational studies; C.L.H., A.R.D., R.P., J.C.B., and C.A.L. provided funding; C.L.H., L.S.Z., A.R.D., R.P., J.H.C., M.A.L., C.A.L. and J.C.B. contributed to methodology; C.L.H. wrote the manuscript; C.L.H., A.R.D., R.P., L.S.Z., and J.C.B. reviewed and edited the manuscript.

## Data and materials availability

Data, MATLAB code, plasmids, and *Toxoplasma gondii* strains are available upon request. Code will be made available on Github by time of publication, and the link will be inserted here on bioRxiv.

