## Supplemental Information for "Emergent Actin Flows Explain Diverse Parasite Gliding Modes"

### Supplementary Information for
### **Emergent Actin Flows Explain Diverse Parasite Gliding Modes**

**Christina L. Hueschen[1,*], Li-av Segev Zarko[2], Jian-Hua Chen[3,4], Mark A. LeGros[3,4,5], Carolyn A. Larabell[3,4,5], John C. Boothroyd[2], Rob Phillips[6,7], and Alexander R. Dunn[1,*]**

[1]Department of Chemical Engineering, [2]Department of Microbiology and Immunology, Stanford University, Palo Alto, California.

[3]Department of Anatomy, University of California San Francisco, San Francisco, California.

[4]National Center for X-ray Tomography, [5]Molecular Biophysics and Integrated Bioimaging Division, Lawrence Berkeley National Laboratory, Berkeley, California.

[6]Department of Physics, [7]Division of Biology and Biological Engineering, California Institute of Technology, Pasadena, California.

**2 Video Legends** **8**

**3 Supplementary Table 1: Observations of Back-and-Forth Gliding** **9**

**4 Experimental Methods and Materials** **10**

**References** **32**

#### 1. Supplementary Figures For Main Text

We begin by presenting, for easy access, the supplementary figures referenced within the body of the main text. Additional figures are embedded through the Supplementary Information document, adjacent to the material that they illustrate or support.

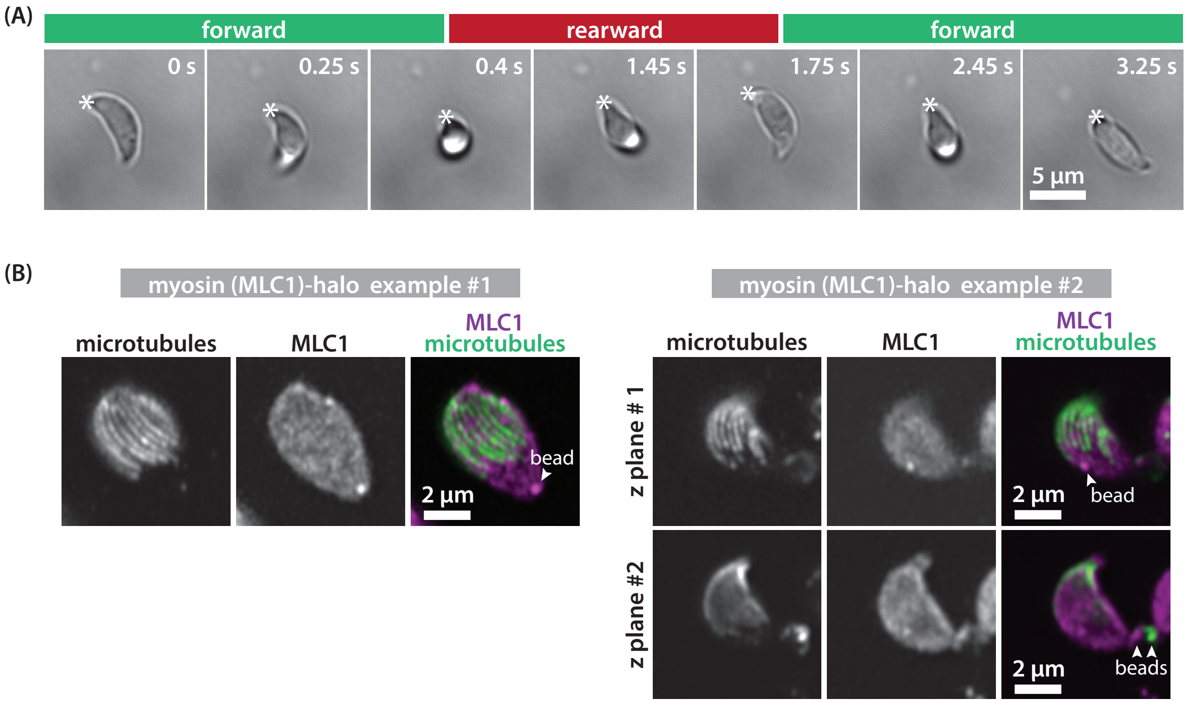

**Figure S1. Back-and-forth patch gliding and MLC1 localization.** (A) Timelapse showing back-and-forth patch gliding behavior in *T. gondii* tachyzoites (parental strain; see Experimental Methods and Materials). Star marks parasite posterior end. (B) Confocal (Airyscan super-resolution) images of fixed extracellular tachyzoites with retracted conoids (left) and protruded conoids (right) expressing myosin light chain 1 (MLC1)-halo and immunostained for $\alpha$-tubulin. MLC1, which recruits myosin A to the IMC [1], does not localize exclusively along subpellicular microtubules. Images shown are individual z-slices. Multicolor beads were used for channel registration.

Figure S1 presents observations inconsistent with a templated model of *Toxoplasma gondii* F-actin transport during gliding, in which F-actin is transported uniformly rearward towards the posterior end of the cell by myosin A. In this model, reviewed in reference [2], a polarized rearward F-actin flow leads to forward traction force and forward (anterior-end-leading) movement of the cell. As highlighted in Figure S1A, however, *Toxoplasma gondii* tachyzoites can move both forward and rearward. Further, rearward F-actin transport was thought to be likely directed or templated by the polarized subpellicular microtubules that lie beneath the inner membrane complex (IMC) [3, 2]. However, as shown in Figure S1B and in references [4, 5], the myosin A light chain MLC1 localizes throughout the IMC, not specifically at subpellicular microtubules. Thus, it remains unclear how the polarity of the microtubules could dictate or restrict the polarity of myosin A activity - and direct an only-rearward F-actin flow.

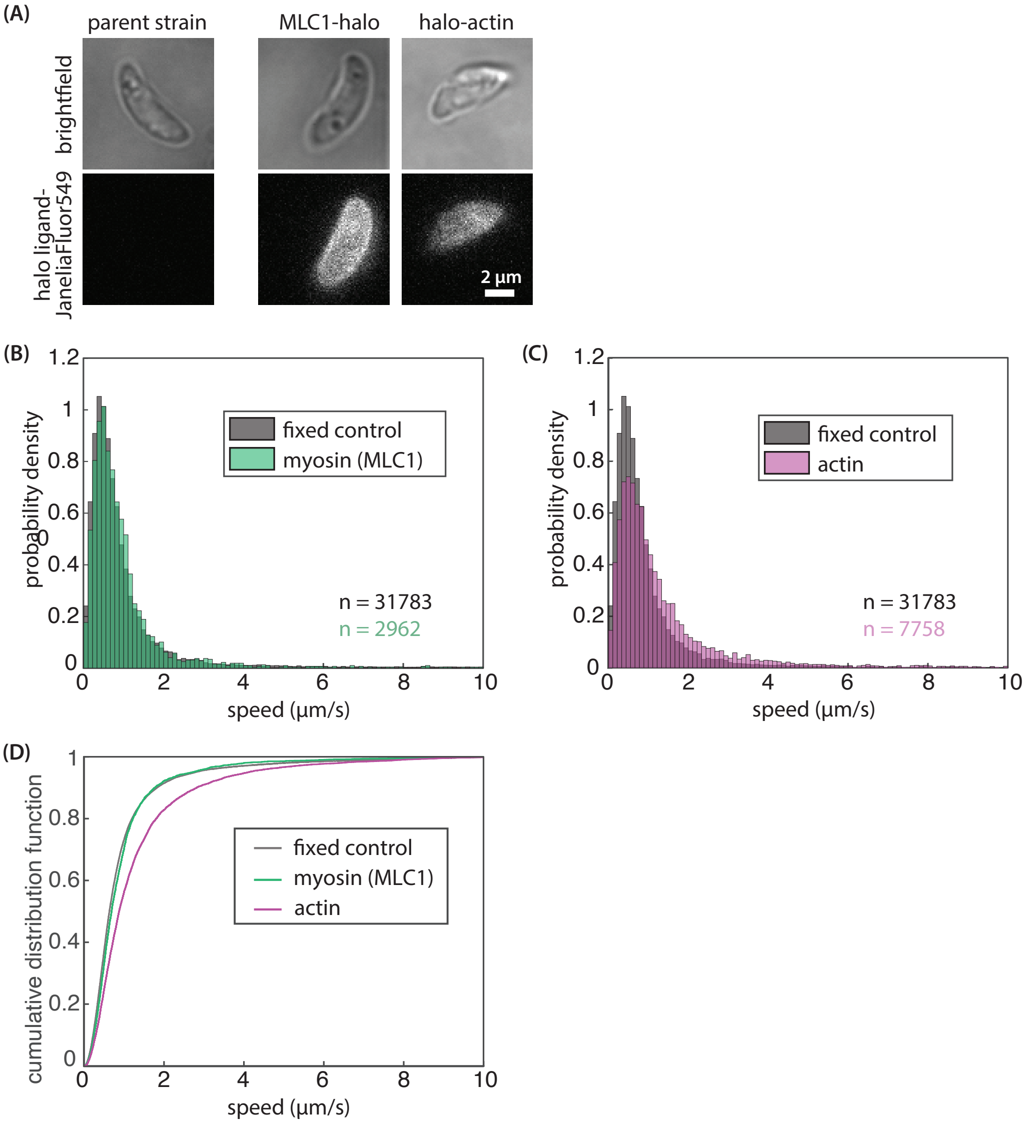

**Figure S2. Imaging and tracking myosin (MLC1) and actin.** (A) Labeling control showing that even at the highest concentrations (500 pM) of HaloTag Ligand Janelia Fluor used, fluorophores specifically labeled only HaloTag fusion proteins. No fluorescent molecules were detectable in parent strain tachyzoites following labeling and standard washes (see Experimental Methods and Materials). (B) Histogram of speeds of automatically-tracked single myosin light chain 1 (MLC1) proteins compared to fixed-cell control. (C) Histogram of speeds of automatically-tracked single actin proteins compared to the same fixed-cell control. In the fixed-cell control, halo-actin proteins are labeled and imaged under identical conditions, but cells are fixed before imaging. This condition allows us to establish the baseline speed distribution for static molecules that arises from measurement error when imaging at the rapid temporal resolution required to capture fast dynamics [6]. Given a pixel size of 100 nm and a frame interval of 86 ms, even a one-pixel localization error produces a measured speed of 1.2 $\mu$m/s. (D) Cumulative distribution function of fixed cell control, MLC1, and actin speeds. (B-D) Automatic tracking reveals a population of mobile actin molecules, compared to MLC1 and fixed control speed profiles (n = 15 cells for actin, 7 cells for myosin, and 6 cells for the fixed control). We note that the actin mobile population is likely an undercount; automatic tracking disproportionately reports immobile and slow trajectories, in which detection events are easier to confidently link. Fast, directional movements visible by eye were challenging to capture with automated methods, necessitating the use of manual tracking for a focused analysis of directional trajectories (Figure 1; as discussed in Supplemental Information Section 5).

Figure S2 presents additional results relevant to the imaging of single myosin light chain 1 (MLC1) and actin molecules inside living extracellular *Toxoplasma gondii* tachyzoites. In these experiments, fusion proteins expressed at low levels (MLC1-halo and halo-actin, see Experimental Methods and Materials) were labeled sparsely with bright, photostable Janelia Fluor dyes using the HaloTag system. The labeling control shown in Figure S2A shows the specificity of this labeling approach. Figure S2B-D reports speeds from automated detection and tracking of MLC1 and actin molecules using the u-track algorithm [7]. While the tracking settings (see Section 5) required to prevent false linkages between molecules disproportionately reports immobile and slow trajectories, in which detection events can be more confidently linked, automated tracking still showed a larger fraction of mobile actin (Figure S2C-D) compared to myosin (MLC1).To put numbers on the difference between the distribution of actin speeds and the distribution of fixed control and myosin speeds visible by eye in FigureS2D, we can calculate the two-sample Kolmogorov-Smirnov test statistic, which reports on the maximum absolute difference between cumulative distribution functions. The Kolmogorov-Smirnov statistic is 0.048 when comparing the speed distributions of immobile fixed-cell control molecules to myosin; 0.15 when comparing the immobile fixed-cell control to actin, and 0.18 when comparing myosin to actin ($p < 0.001$ in all 3 cases, suggesting that each distribution is unique). Importantly, we note that the challenge of using automated methods to capture fast, directional actin movements visible by eye necessitated the use of manual tracking for a focused analysis of directional trajectories shown in Figure 1 D-F (see Section 5 for additional discussion).

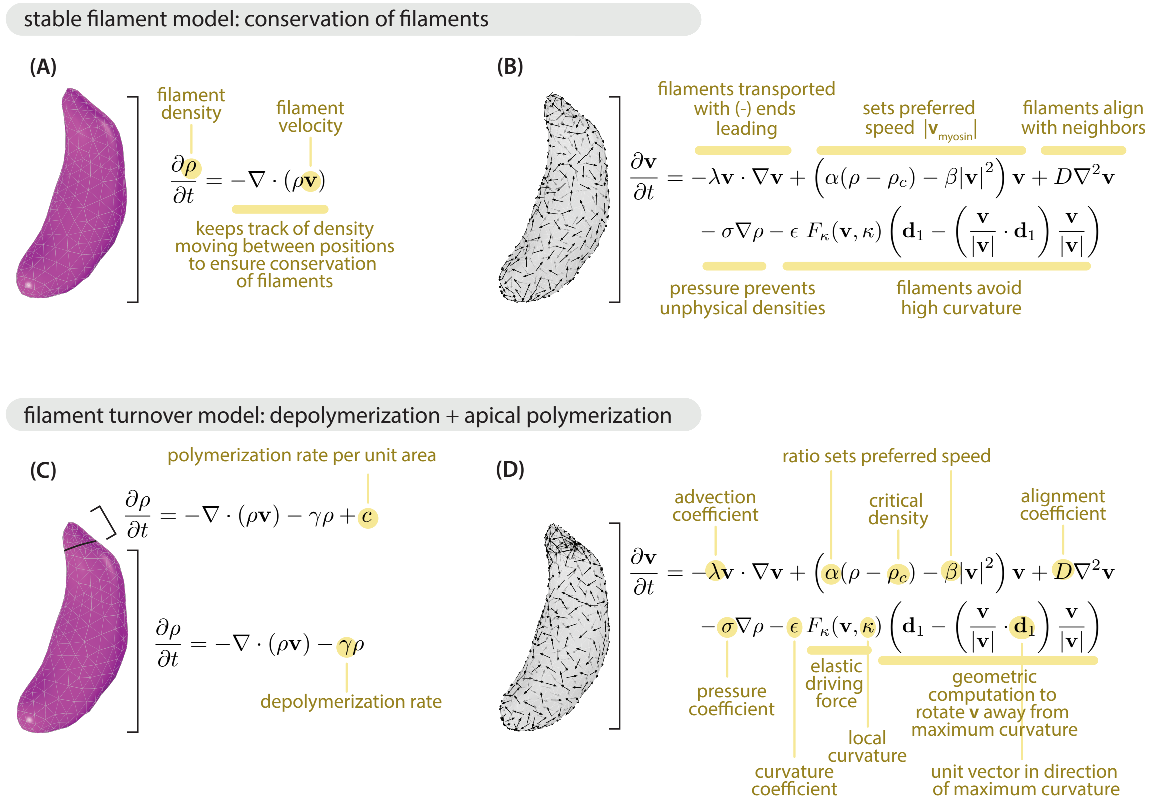

**Figure S3. Governing equations of actin filament self-organization.** (A) In a regime in which filaments are stable and conserved, the continuity equation governs filament density. (B) Generalized Toner-Tu flocking theory for actin filament self-organization. Different terms correspond to different effects governing filament velocity and speed. In other words, filament velocity evolves according to local rules, implemented mathematically. (C) In a filament turnover regime, incorporating our knowledge of *Toxoplasma* actin biology, we allow filament polymerization at the cell's anterior (apical) end and filament depolymerization throughout. (D) Filament velocity evolves according to the same equation (B) as in the stable filament regime. We include this second version in order to pedagogically annotate the coefficients that tune each term.

In Figure S3, we annotate the Toner-Tu flocking model used to predict emergent actin filament organization and dynamics at the *Toxoplasma gondii* cell surface. For the interested reader, the model is presented in detail in Section 6 and reference [8]. In brief, actin filament organization is described by two fields: the scalar field $\rho$, which captures filament density, and the velocity vector field $\mathbf{v}$, which captures both filament polarity (orientation of v) and speed (magnitude of v). The continuity equation ensures conservation of stable filaments (Figure S3A) or is modified to capture filament polymerization at the cell anterior end and filament depolymerization throughout (Figure S3C). As shown in Figure S3B and D, filament velocity evolves according to a minimal version of the Toner-Tu flocking model, with an additional term that penalizes filament curvature which is derived in Section 6.3.

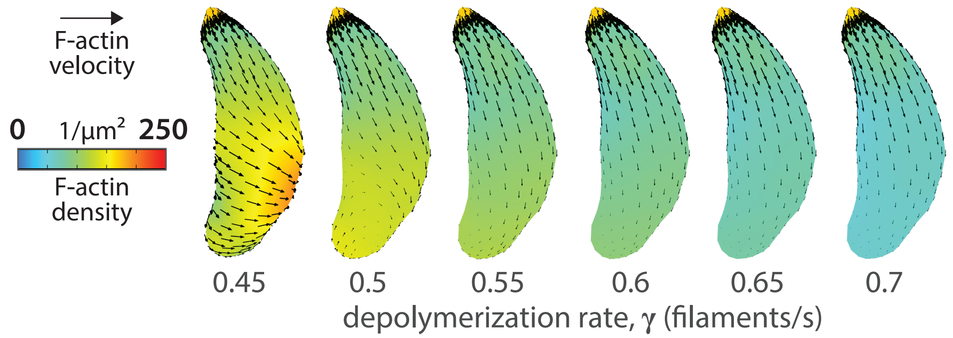

**Figure S4. Predicted steady-state F-actin density and velocity patterns for different filament turnover rates.** Tuning actin depolymerization rate ($\gamma$, filaments/s) in the Toner-Tu actin self-organization model changes features like chirality and density gradient in the emergent steady-state actin patterns predicted. Shown here are steady-state filament density (color) and velocity (black arrows) for a filament polymerization rate $c = 1500\ \mu m^{-2} s^{-1}$ (estimated in Section 7.2) and filament turnover rates $\gamma$ as indicated.

In Figure S4, we highlight the effect of actin filament depolymerization rate on predicted steady-state unidirectional F-actin flows. These flow patterns, which we call "steady-state" because the density and velocity fields reach a stable solution which they hold over time, consist of largely unidirectional, rearward actin flow and are labeled as "unidirectional" (dark purple circles) in Figure 4B. Those steady states that lie in a regime close to the transition to recirculation are most consistent with cell motions during helical gliding, circular gliding, and twirling, as depicted in Figure 4D-F. In the example shown here, this regime corresponds to depolymerization rates of $\gamma$ = 0.45-0.55 filaments/s. (Lower rates of depolymerization lead to F-actin recirculation ("cyclosis," orange circles), as shown in Figure 4B.) Our theoretical model indicates that in *Toxoplasma gondii* parasites whose actin dynamics lie in this regime, actin self-organization alone is sufficient to generate patterns that explain helical gliding, circular gliding, and twirling. Interesting, this finding is consistent with experimental results suggesting that careful tuning of actin filament stability is critical for normal gliding behaviors [9, 10, 11, 12] We note that this sufficiency does not preclude the existence of additional complexity not explored here, like contributions from inner membrane complex (IMC) shape.

#### 2. Video Legends

**Video 1. Diverse gliding modes: an individual cell performs back-and-forth gliding, helical gliding, and then twirling.** In this brightfield microscopy video, an extracellular *Toxoplasma gondii* tachyzoite glides back and forth (0:00 min:s), displaying so-called patch or pendulum gliding (see Table S1). Several minutes later, the same cell displays helical gliding (4:04 min:s), followed by twirling (4:13 min:s). The ability of an individual cell to switch between gliding modes on the timescale of minutes is consistent with the self-organization hypothesis presented in this work. Different self-organized actin states may arise at different points in time, likely in response to altered regulation of actin dynamics, even though the underlying cell structure (e.g., IMC and microtubules) remains unchanged.

**Video 2. Single-molecule imaging reports on dynamics of myosin in extracellular *Toxoplasma gondii.*** Example of total internal reflection fluorescence (TIRF) imaging of myosin light chain 1 (MLC1-halo) single molecule dynamics (cyan) in an extracellular parasite, with bulk labeling (magenta) to show cell position. MLC1-halo was expressed at low levels, labeled with picomolar concentrations of Janelia Fluor 549 to visualize single molecules (cyan), and labeled with nanomolar concentrations of Janelia Fluor 647 to visualize the cell (magenta). MLC1 molecules frequently remained immobile, or bound, on the time scale of seconds. Five molecules, indicated by arrowheads, remain bound for the length of the movie. Time is in min:s.

**Video 3. Single-molecule imaging reports on dynamics of actin in extracellular *Toxoplasma gondii.*** Example of total internal reflection fluorescence (TIRF) imaging of *Toxoplasma* actin (halo-actin) single molecule dynamics (cyan) in an extracellular parasite, with bulk labeling (magenta) to show cell position. Halo-actin was expressed at low levels, labeled with picomolar concentrations of Janelia Fluor 549 to visualize single molecules (cyan), and labeled with nanomolar concentrations of Janelia Fluor 647 to visualize the cell (magenta). The same video repeats three times in order to highlight different molecule behaviors. First, "b" labels an immobile ("bound") molecule, which persists for a second before disappearing - likely because it unbinds and moves out of the TIRF field. Second, "d" labels a molecule that displays meandering (diffusive) behavior. Third, "m" labels a directional ("motor-transported") molecule that moves persistently towards the anterior of the cell. Time is in min:s.

**Video 4. Actin flocking model predicts self-organized recirculation ("cyclosis") of actin patches in the absence of filament turnover.** The simulation begins with a disordered network, and then filament density $\rho$ and velocity $\mathbf{v}$ evolve over time on the *Toxoplasma gondii* tachyzoite cell surface according the actin self-organization Toner-Tu equations presented in Sections 6 - 9. Filaments are stable and conserved, as in Supplementary Figure S3 A-B. Equations were solved and results simulated using the finite element method in COMSOL Multiphysics®. The cell surface shape was obtained by soft X-ray tomography.

**Video 5. Imaging of jasplakinolide-stabilized actin bundles that recirculate up and down the *Toxoplasma gondii* cell.** Example of total internal reflection fluorescence (TIRF) imaging of halo-actin in extracellular *Toxoplasma* tachyzoites treated briefly with 1 $\mu M$ jasplakinolide to stabilize actin filaments. The recirculating bundle sometimes moves out of the TIRF field but is clearly visible when parallel to the imaging plane (e.g., 0:13-0:15 min:s).

**Video 6. Actin flocking model predicts the emergence of self-organized unidirectional flow in the presence of filament turnover.** The simulation begins with a disordered network, and then filament density $\rho$ and velocity $\mathbf{v}$ evolve over time on the *Toxoplasma gondii* tachyzoite cell surface according the actin self-organization Toner-Tu equations presented in Sections 6 - 9. Filaments are polymerized in the conoid (anterior end) at rate $c = 1500\ \mu m^{-2} s^{-1}$ and depolymerized throughout the cell surface at rate $\gamma\rho = (0.5\ s^{-1})\ (\rho\ \mu m^{-2})$, as in Supplementary Figure S3 C-D and Section 7.2. Equations were solved and results simulated using the finite element method in COMSOL Multiphysics®. The cell surface shape was obtained by soft X-ray tomography.

##### 3. Supplementary Table 1: Observations of Back-and-Forth Gliding

| Organism & life stage | Condition | Expected effect on actin | Observed bidirectional gliding phenotype | Reference |
|---|---|---|---|---|
| *T. gondii* tachyzoites | "Wild-type" extracellular RH in media conditioned by human foreskin fibroblasts | N/A | "Patch" gliding with frequent pauses: e.g., gliding forward, rearward, forward, flipping over the posterior cell end, rearward, pausing, etc. Cell-substrate contact point is stationary in the lab reference frame. | This study (Fig. S1, Video 1) |
| *T. gondii* tachyzoites | Jasplakinolide treatment | Increased filament stability | "frequent reversal of direction often resulted in a behavior called "rolling," during which the parasite moved forward, raising its anterior end off the substrate, then reversed course, elevating the posterior end." | Wetzel et al., 2003 [9] |
| *T. gondii* tachyzoites | Small molecule treatment | Unknown targets | "parasites... move quickly with frequent changes in direction" | Carey et al., 2004 [13] |
| *T. gondii tachyzoites* | "ADF cKO" (actin depolymerizing factor conditional knockout) | Increased filament stability | "frequent reversals of direction leading to no net movement... back-and-forth rocking motions" | Mehta & Sibley, 2011 [10] |
| *T. gondii* tachyzoites | "DOC2.1 mutation" | Blocked microneme secretion | "shuffling, a distinct motility mode in which parasites abruptly move back and forward" | Farrell, Thirugnanam et al., 2012 [14] |
| *P. berghei* sporozoites | "coronin(-)" | Loss of filament crosslinking; possibly increased filament stability due to decreased ADF recruitment | "often detached from the substrate, frequently moving just back and forth over a single adhesion site... In addition... bending and flexing movements without moving forward" | Bane et al., 2016 [15] |
| *P. berghei* sporozoites (from hemolymph) | "Wild-type" sporozoites | N/A | "Patch" gliding: "sporozoites continuously move over a single spot in a back-and-forth manner at similar speeds in both directions" | Münter et al., 2009 [16] |

| Organism & life stage | Condition | Expected effect on actin | Observed bidirectional gliding phenotype | Reference |
|---|---|---|---|---|
| *P. berghei* sporozoites (from hemolymph) | "trap(-)" (truncated adhesin protein TRAP [17]) isolated from mosquito hemolymph | Unknown; truncated TRAP may fail to bind an adaptor (GAC homolog?) that regulates F-actin stability[18] | "Patch" gliding: "sporozoites continuously move over a single spot in a back-and-forth manner at similar speeds in both directions" | Münter et al., 2009 [16] |
| *P. berghei sporozoites* (from midgut) | Mutations in C-terminal cytoplasmic tail of TRAP | Unknown; mutants fail to bind an adaptor (GAC homolog?) that regulates F-actin stability [18] | "Pendulum" gliding: "repeated cycles of (a) gliding over one third of a circle, (b) stopping for usually 1-2 s, and (c) moving back to the original position" | Kappe et al., 1999 [19] |
| *P. berghei* sporozoites (from salivary gland) | Mutations in actin subdomain 4 | Reduced actin filament turnover | "parasites... frequently paused and reversed direction during migration" | Douglas et al., 2018 [11] |
| *P. berghei* sporozoites (from midgut, hemolymph) | "cb$\beta$ (-)": capping protein subunit b loss of function | Unrestricted filament polymerization | "cp$\beta$(-) sporozoites display only non-productive motility patterns, such as bending, flexing and pendulum movement" | Ganter et al., 2009 [12] |

#### 4. Experimental Methods and Materials

##### *4.1. Parasite and host cell culture*

*Toxoplasma gondii* tachyzoites were maintained by serial passage in primary human foreskin fibroblasts (HFFs) in Dulbeccos modified Eagles high glucose medium (DMEM; Gibco 11960-044) with 10% heat-inactivated fetal bovine serum (FBS; Corning 35-011-CV), 2 mM glutamine (Sigma-Aldrich G7513), 100 U/ml penicillin, and 100 $\mu$g/ml streptomycin (Gibco 15140122) at 37°C in 5% $CO_2$. In brief, to passage parasites, infected HFF monolayers were suspended in media by scraping, syringe lysed using a 25-gauge blunt-end needle (SAI Technologies B25-50), and added at a 150-fold dilution to an confluent uninfected HFF monolayer, every 2-4 days. HFFs were obtained from the neonatal clinic at Stanford University following routine circumcisions that are performed at the request of the parents for cultural, health, or other personal medical reasons (i.e., not in any way related to research). These foreskins, which would otherwise be discarded, were fully deidentified and therefore do not constitute human subjects research. Uninfected HFFs were maintained in the supplemented DMEM described above, passaged using 0.25% trypsin-EDTA (Gibco 25200056), and discarded after passage 15.

##### *4.2. Generation of halo-ACT1 and MLC1-halo strains*

In brief, halo-TgACT1 or TgMLC1-halo fusions under the control of a weak promoter (from TGGT1_239010, gift of M. Panas [20]) were incorporated into the genome of the *Toxoplasma gondii* type I RH hxgprt ku80 strain [21]. In detail, p239010-halo-C1-HXGPRT and p239010-halo-N1-HXGPRT vectors were created by replacing a region of the pGRA-3xHA-HPT vector [22] (from the pGRA promoter through the translated region) with: (1) the TGGT1_239010 promoter and 5' UTR to drive low expression levels; (2) the HaloTag sequence (Promega); and (3) a serine-glycine linker and multiple cloning site either C-terminal ("C1") or N-terminal ("N1") to the HaloTag sequence. TgACT1 (TGGT1_209030) or TgMLC1 (TGGT1_257680) were then synthesized and cloned into the p239010-halo-C1-HXGPRT or p239010-halo-N1-HXGPRT vector, respectively (Epoch Life Science, Inc., Missouri City, TX). For transfection with p239010-halo-ACT1-HXGPRT or p239010-MLC1-halo-HXGPRT, RH Δhxgprt Δku80 parasites were mechanically released in PBS, pelleted, and resuspended in 20 μl P3 primary cell Nucleofector solution (Lonza) with 15 μg DNA, and electroporated using the Amaxa 4D Nucleofector (Lonza). Transfected parasites were permitted to infect and grow in confluent HFFs for 48 h, after which time the media was supplemented with 50 μg/ml mycophenolic acid and 50 μg/ml xanthine for HXGPRT selection. Parasites were passaged 4 times over 10-12 days in selection media before being singly cloned into 96-well plates by limiting dilution. Clones were expanded and screened for HaloTag expression by incubating intracellular parasites overnight with 50 nM TMR HaloTag Ligand (Promega G8251), washing 5x with PBS to remove unbound dye, fixing with 4% paraformaldehyde (EMS AA433689M) for 15 min, and imaging fluorescence.

##### *4.3. Single molecule labeling in live parasites*

Before infected HFFs were lysed to release parasites for imaging, parasites within infected HFF monolayers were labeled for 3 h at 37°C with Janelia Fluor 549 HaloTag Ligand (Promega GA1110) and Janelia Fluor 646 HaloTag Ligand (Promega GA1120) [23] at a concentration of 1-10 pM for single molecule imaging or 100-500 pM for bulk population imaging. Because Janelia Fluor dyes bleach over time in storage, dye concentration must be optimized empirically and adjusted on the timescale of months; furthermore, care should be taken to avoid more than two or three freeze-thaw cycles before use. When subpellicular microtubule imaging was used to determine parasite polarity, infected HFFs were labeled with 100 nM siR-tubulin and 10 μM verapamil (Cytoskeleton, Inc. CY-SC002) alongside 1-10 pM Janelia Fluor 549. Before parasite release, infected HFF monolayers were washed 7x with DMEM to ensure removal of unbound dye.

##### *4.4. Preparation and TIRF imaging of live extracellular parasites*

To release parasites, infected HFFs were scraped and syringe lysed in fresh phenol red-free DMEM with a 27-gauge needle (SAI Technologies B27-50). Freshly released parasites were placed on 35 mm #1.5 glass-bottomed dishes (Cellvis D35-20-1.5-N; incubated with 10% FBS before use) with a confluent monolayer of human foreskin fibroblasts (HFFs) grown on Snapwell Insert polyester membranes (Corning Costar CLS3801) suspended approximately 0.2 mm above them. Parasites were imaged at 30°C using objective-type total internal reflection fluorescence (TIRF) microscopy on an inverted microscope (Nikon TiE) with a heated Apo TIRF 100 oil objective of numerical aperture 1.49 (Nikon) and controlled using Micromanager [24]. To enable simultaneous two-color imaging, samples were excited with both 532 nm (Crystalaser) and 635 nm (Blue Sky Research) lasers, and emitted light passed through a quad-edge laser-flat dichroic with center/bandwidths of 405 nm/60 nm, 488 nm/100 nm, 532 nm/100 nm, and 635 nm/100 nm from Semrock (Di01-R405/488/532/635-25x36) and corresponding quad-pass filter with center/bandwidths of 446 nm/30 nm, 510 nm/30 nm, 581 nm/30 nm, 703 nm/30 nm band-pass filter (FF01- 446/510/581/703-25). Emission channels were then separated as previously described [25] and recorded on an electron-multiplying charge-coupled device (EMCCD) camera (Andor iXon).

##### *4.5. Jasplakinolide treatment and results*

Live extracellular parasites were prepared as in subsection 4.4, with the addition of 1 μM jasplakinolide (Millipore Sigma J4580) immediately before imaging. In a narrow window of time from approximately 15-30 min after jasplakinolide addition, protruding bundles of actin filaments were observed circling around the periphery of parasites. We note that this recirculating behavior was very sensitive to treatment time and drug concentration. Over time, most protrusions lost this recirculating behavior and became fixed in position at the anterior (apical) end, as previously observed [26]. We speculate that this transition to fixed apical actin bundles occurs as bundles grow long enough (with polymerization favored both by jasplakinolide and by apically-localized FRM1 [18]) to protrude through the conoid and into the cytoplasm and can no longer re-orient to contact myosin motors on the outside of the inner membrane complex. Under ideal treatment conditions, nearly all extracellular parasites observed displayed recirculating actin protrusions (n ≈ 30 cells); under less ideal conditions (e.g., after more than 30 min treatment, or with poorly attached parasites), less than 10% of parasites displayed recirculating protrusions (n ≈ 50 cells).

##### *4.6. MLC1 immunofluorescence and super-resolution confocal microscopy*

Parasites were released from infected HFF monolayers by scraping and syringe lysis in DMEM with a 27-gauge needle (SAI Technologies B27-50), passed through a 5 $\mu$m filter (Millipore Sigma SLSV025LS) and allowed to settle onto #1.5 coverslips at 37°C in 5% $CO_2$ for 30 min in DMEM + 1 $\mu$M calcium ionophore A23187 (Sigma C7522). Subsequent staining steps were performed at room temperature: parasites were fixed with warm 4% paraformaldehyde (EMS AA433689M) for 15 min, washed 3x with phosphate-buffered saline (PBS), incubated with permeabilization and blocking buffer (PBS-T + 2% BSA = 0.1% Triton-X-100 and 2% bovine serum albumin in PBS) for 20 min, incubated with mouse anti--tubulin monoclonal antibody DM1$\alpha$ (Sigma T6199; diluted 1:500) and 2 nM Janelia Fluor 646 HaloTag Ligand (Promega GA1120) in PBS-T + 2% BSA for 1 h, washed 3x with PBS, incubated with anti-mouse IgG secondary antibody conjugated to Alexa Fluor 488 (Cell Signaling 4408S) for 20 min, washed 3x with PBS, and mounted in ProLong Gold Antifade (ThermoFisher P36934). Samples were imaged using an inverted Zeiss LSM 780 confocal microscope with a 63X/1.4 NA oil objective, 488 nm Ar laser, 633 nm HeNe laser, and a Zeiss Airyscan detector (32-channel gallium arsenide phosphide photomultiplier tube (GaAsP-PMT) area detector), in which using each detector element as an individual pinhole combined with linear deconvolution achieves a spatial resolution below the diffraction limit [27]. All images were acquired using Zen Black software (Carl Zeiss).

##### *4.7. Soft X-ray tomography*

HFF monolayers, 18-20 hours after parasite infection, were washed two times with Hanks balanced salt solution (HBSS; Gibco 14175095) supplemented with 1 mM magnesium chloride, 1 mM calcium chloride, 10 mM sodium hydrogen carbonate and 20 mM HEPES, pH 7. HFFs were scraped and passed through a 27-gauge needle (SAI Technologies B27-50) to release parasites into fresh HBSS at room temperature. Calcium ionophore A23187 (Sigma C7522) at a final concentration of 1 $\mu$M was added to the sample at room temperature for 10 minutes. Parasites were pelleted, access liquid was aspirated, and parasites were resuspended in the remaining liquid (≈25 $\mu$l) prior to loading into 5 $\mu$m-diameter glass capillaries. Parasites inside capillaries were then vitrified by fast plunge-freezing in 90 K liquid propane. Capillaries were imaged using the XM-2 cryo soft X-ray microscope (SXM) at the National Center for X-Ray Tomography at the Advanced Light Source (Lawrence Berkeley Laboratories, Berkeley, CA). The XM-2 is equipped with a micro zone plate (MZP) with a spatial resolution of 60 nm, and the imaged capillary was in an atmosphere of helium gas stream cooled by liquid nitrogen. To have a full rotated tomographic dataset reconstructed, 92 projection images were taken with 2 degree increments. The exposure time of each projection varied between 200-450 ms, depending on the beam flux and the sample thickness. Projection images were normalized and aligned, and the tomographic reconstructions were calculated using iterative reconstruction methods in the AREC-3D package [28]. Additional information on the soft X-ray tomography method is available within reference [29].

#### 5. Image Analysis and Single Molecule Tracking

Automated molecule detection and tracking (Supplementary Figure S2) was performed on raw movies of halo-actin and MLC1-halo molecules in live parasites using u-track software (release 2.2.0) made available by the Danuser lab [7] and run through MATLAB R2019a from Mathworks, Inc. Key u-track parameters were as follows: $\alpha = 0.05$ (sets spot detection threshold relative to background); Gaussian standard deviation $\sigma = 1.3$ pixels = 130 nm (based on an average measured full width half maximum (FWHM) of 3 pixels for single spots, and the relationship between FWHM and $\sigma$ for the Gaussian distribution: FWHM $= 2\sqrt{2\ ln2}\ \sigma \approx 2.355\ \sigma$); search radius = 0-9 pixels, corresponding to a maximum allowed speed of 9 pixels / 86 ms = 10.5 $\mu m\ s^{-1}$; gap tolerance = 0; minimum track length = 5 frames = 0.43 s. These u-track settings were selected empirically to minimize false positive detection events and linkages (erroneous inclusion of two different molecules in a single trajectory), but they necessarily under-report fast-moving population trajectories. After much discussion and consideration of alternative tracking approaches and algorithms, we came to the realization that every automated algorithm is benchmarked against the "gold standard" of object tracking by the human brain, and that manual human tracking of the fast directional actin trajectories apparent by eye was not only a valid option, but in fact the most accurate one available. Thus, the fast, directional actin population tracks (Figure 1D-F) were obtained by manual spot tracking of raw images with the Manual Tracking plugin [30] within Fiji (ImageJ Version 2.0.0-rc-69/1.52p) [31].

Single molecule trajectory outputs (position over time) from u-track or manual tracking were analyzed using home-written MATLAB programs. To align molecule trajectories from different cells to a "reference cell" and calculate the orientation of molecule velocities with respect to cell polarity, cell orientation was determined and trajectories were rotated as follows. First, cell anterior-posterior polarity was determined for every movie analyzed using either subpellicular microtubule imaging or by tracking the cell posterior end following twirling events, in which the posterior end can be identified as the end in contact with the substrate. We note that posterior nuclear position proved to be a consistent polarity indicator as well; while we did not consider nuclear position alone sufficient information to include a cell in this polarity analysis, polarity information from nuclear position analysis was always in agreement with microtubule imaging or twirling analysis. Second, within a home-written MATLAB program, cells were segmented using a threshold determined by Otsu's method (MATLAB Image Processing Toolbox) and fit with an ellipse (regionprops; MATLAB Image Processing Toolbox). Ellipse centroid and orientation (combined with manually annotated polarity info, giving the ellipse an anterior-posterior polarity) were used to determine a rotation matrix that, when applied to cell images or molecule trajectories, aligned them to a "reference cell" with anterior (apical) end pointing up.

All analyses described above were performed on raw images. For display in figures and movies, images were denoised using noise2void [32].

#### 6. Theoretical Model of *Toxoplasma gondii* Actin Filament Self-Organization

##### *6.1. Continuum flocking theory: background and model choice*

To describe the collective motion and organization of actin filaments, we repurposed a classic continuum active matter model that was originally developed by John Toner and Yuhai Tu, inspired by the work of Támas Vicsek, to describe the collective behavior of flocking or schooling animals [33, 34, 35]. This class of theoretical models, known as Toner-Tu or flocking theory, describe collections of "dry," polar, self-propelled agents at any lengthscale, from flocks of flying birds to collections of polarized cytoskeletal filaments. In our case, *Toxoplasma gondii* actin filaments at the cell surface are propelled along by an underlying carpet of plus-end-directed myosin motors, whose action we can effectively capture as polarized filament self-propulsion. One final consideration for us, in choosing a useful and appropriate continuum model for our system, was the distinction between so-called "wet" and "dry" active matter models. In wet active matter systems, the total momentum of self-propelled agents and the media they live in is conserved. Dry active matter are systems without momentum conservation, often due to dominant frictional drag during movement along a surface, or between two walls [36]. In our system of interest, actin filaments move through a confined space $\approx 25$ nm in height between the *T. gondii* plasma membrane and the rigid, intermediate filament-reinforced inner membrane complex [1], which we expect to impose a no-slip boundary condition. To make sure that modeling our *T. gondii* actin filament network as a dry, polar system (using Toner-Tu flocking theory) is indeed justified, we now turn to estimating expected viscous drag and friction scales. In the remainder of this subsection, we present this estimate and find that frictional drag indeed dominates. In the subsection that follows, we move on to the details of the Toner-Tu model and its adaptation for our purposes.

One way to formalize the distinction between wet and dry active matter is in terms of a hydrodynamic length scale,

$$l = \sqrt{\frac{\eta}{\gamma}} \tag{1}$$

which reflects a competition between the viscosity $\eta$ and a friction coefficient $\gamma$. In dry active matter systems dominated by frictional drag, $l$ is very small. In a sense, hydrodynamic "communication" through the intervening fluid medium is only possible over very short distances and can thus be neglected.

In our case, actin filament dynamics play out within an extremely thin layer ($\approx 25$ nm thick) above a no-slip boundary. We consider the magnitude of viscosity and of friction for the intervening fluid (cytoplasm) surrounding our actin filaments. We take $10^{-3}\,\mathrm{N{\cdot}s/m^2}$, the viscosity of water, as a reasonable estimate for $\eta$. Next, we require an estimate for the parameter $\gamma$, given our knowledge of actin filament dynamics and subcellular structure at the *Toxoplasma* cell surface. We present this estimate in Appendix Section 10.1. In brief, by understanding in-plane frictional drag as capturing viscous interactions across the height of a thin three-dimensional flow, we estimate a friction coefficient $\gamma \approx 10^{13}\,\mathrm{N \cdot s/m^4}$.

Following eqn. 1, we estimate a hydrodynamic length scale

$$l \approx \sqrt{\frac{10^{-3}\,\frac{\mathrm{N\cdot s}}{\mathrm{m^2}}}{10^{13}\,\frac{\mathrm{N\cdot s}}{\mathrm{m^4}}}} \approx 10^{-8}\,\mathrm{m} \tag{2}$$

or 10 nm, a distance smaller than the scale of our agents (filaments) themselves, or their estimated spacing. Thus, we choose a dry, polar active matter model (Toner-Tu flocking theory) to describe the collective motion and self-organization of actin filaments at the parasite surface.

##### *6.2. A minimal Toner-Tu flocking theory*

The continuum flocking equations developed by Toner and Tu use two field variables to characterize the evolution of a flock (a collection of birds, or sheep, or actin filaments) in space and time: density $\rho(\mathbf{r}, t)$ and velocity $\mathbf{v}(\mathbf{r}, t)$. Rather than keeping track of each discrete agent (every bird, sheep, or filament) over time, we use the powerful approach of coarse-graining. In our case, this means "dividing up" the *Toxoplasma gondii* cell surface into a continuous field of boxes, or area elements, and identifying an average actin filament velocity and density for each box. This continuum approach enables us to predict the time-evolution and emergence of cell-scale actin filament patterns in the language of our field variables, filament density $\rho$ and velocity $\mathbf{v}$. Mathematically, the evolution of our two field variables in space and time is carried out in the language of partial differential equations. We can think of these partial differential equations as "update rules" that tell us how to use our knowledge of the field variables $\rho(\mathbf{r}, t)$ and $\mathbf{v}(\mathbf{r}, t)$ at a time $t$ and work out their subsequent values $\rho(\mathbf{r}, t + \Delta t)$ and $\mathbf{v}(\mathbf{r}, t + \Delta t)$ at time $t + \Delta t$. This update process repeats, one time step after another, to get the full space-time history of filament density and velocity on the cell surface. We note that, in acknowledgement that our actin filaments of interest are confined in a thin layer between the inner membrane complex and the plasma membrane, our density and velocity fields are strictly two-dimensional, confined to the tangent plane of the cell surface.

With field variables in hand, we write the governing field equations (i.e. partial differential equations, or "update rules") that describe their spatiotemporal evolution. The first governing equation is the continuity equation,

$$\frac{\partial \rho}{\partial t} + \nabla \cdot (\rho \mathbf{v}) = 0. \tag{3}$$

Conceptually, all that this equation says is that if we consider an infinitesimal box of material, the change in the amount of actin within that little box is the difference between the amount that flows in and the amount that flows out. For the moment we consider the case of strict mass conservation with no sources or sinks, though we will see below that the biology of *Toxoplasma* requires us to soften that constraint and allow for both polymerization and depolymerization of actin filaments. The second governing equation is a minimal representation of the dynamics of the $\mathbf{v}$ field offered by Toner and Tu and given first in direct notation by

$$\frac{\partial \mathbf{v}}{\partial t} = [\alpha(\rho - \rho_c) - \beta |\mathbf{v}|^2]\mathbf{v} + D\nabla^2 \mathbf{v} - \sigma \nabla \rho - \lambda \mathbf{v} \cdot \nabla \mathbf{v} \tag{4a}$$

and then, equivalently, in index notation by

$$\frac{\partial v_i}{\partial t} = [\alpha(\rho - \rho_c) - \beta v_j v_j] v_i + D\nabla^2 v_i - \sigma \frac{\partial \rho}{\partial x_i} - \lambda v_j \frac{\partial v_i}{\partial x_j}, \tag{4b}$$

where $\rho_c$ is the critical density above which the filaments move coherently, the square root of the ratio of $\alpha(\rho - \rho_c)$ and $\beta$ sets a filament transport speed scale $|\mathbf{v}| = \mathrm{v}_{\mathrm{myosin}}$, the coefficient $D$ tunes filament alignment with neighbors, $\sigma \nabla \rho$ provides an effective pressure that keeps filament density within a physical range, and the coefficient $\lambda$ tunes filament velocity self-advection. Note that we are using the summation convention, which tells us to sum over all repeated indices; for example, $\mathbf{a} \cdot \mathbf{b} = a_i b_i = \sum_{i=1}^{3} a_i b_i$.

As illustrated in Figure 2 and Supplementary Figure S3, these terms are a mathematical encapsulation of the "rules" of local filament interactions with other filaments, with myosin motors, and with the geometry of their environment. Together, they provide an update to filament velocity, over each increment in time. Put differently, each term captures a "driving force" that acts locally on filament velocity. The first term on the right, preferred speed, sets the characteristic speed of the actin filaments at 5 $\mu$m/s, the approximate mean speed of the relevant myosin motor, TgMyoA (Figure 1 and [37]). The second term, neighbor coupling, involving the Laplacian, we can think of as a velocity smoothing term which pushes a given filament's velocity (orientation and speed) to better match that of its neighbors. For the case of *T. gondii* actin filaments, this neighbor alignment term captures the action of the filament cross-linking protein coronin [17, 38, 15] and polar alignment through physical collision of moving filaments, as seen *in vitro* [39]. The third term is a pressure term that punishes gradients in density, keeping filament density within a realistic range.

The final term, velocity advection, has analogy to the gradient component of the material time derivative in the Navier-Stokes equations. In essence, the velocity field advects itself; filaments move along in a direction dictated by their orientation and velocity, and they bring that orientation and velocity with them. In the simple case of pure self-advection, $\lambda = 1$. For active agents, such as birds in a flock, $\lambda$ maybe be further tuned by the behavioral response of the agents to gradients in velocity. In other words, $\lambda = 1 + \xi$, where $\xi$ reflects a behavioral "gradient penalty"; for example, birds may resist flying quickly into a steep gradient of decreasing velocity, and may slow down. For the case of *T. gondii* actin filament self-organization, we choose $\lambda = 1$ as discussed in Section 7.2, because our knowledge of this myosin-driven actin network does not suggest an active ability of filament velocity to respond to nearby gradients in velocity. However, we maintain the advection coefficient $\lambda$ in our general presentation of the theory, for consistency with the Toner-Tu tradition and to provide the most generally useful version of these minimal flocking equations.

Each term of the four terms in these equations is necessary to capture the basic phenomenology of flocking. In the absence of the preferred speed term or the velocity advection term, filaments do not move. In the absence of the neighbor coupling term, they remain disordered, with no collective motion. In the absence of the pressure term, aphysical densities emerge (negative densities, or densities higher than a cystalline packing of filaments).

We hope that the discussion thus far has clearly presented the mathematics and the intuition of a minimal Toner-Tu flocking theory. We refer the interested reader to reference [8] for additional discussion of this theory, its formulation in a general surface form using extrinsic differential geometry, and a numerical exploration of flocking solutions on varied curved surface shapes. Here, we next extend the minimal flocking theory with new terms specific to *Toxoplasma gondii* actin biology and present our full model for actin filament self-organization on the surface of *Toxoplasma*.

##### *6.3. Extending flocking theory to capture Toxoplasma actin biology: curvature*

The original Toner-Tu flocking theory was motivated largely by the motions of animals at macroscopic scales. For the actin filament self-organization in *Toxoplasma*, we introduce several new terms in the velocity and density "update rules." First, we acknowledge that for our actin filaments of interest, confined between the parasite plasma membrane and the inner membrane complex, there is an energetic penalty for filaments whose locally curved environment induces filament bending. Indeed, for actin filaments of approximately 100 nm in length ($l$), with a persistence length ($L_p$) $\approx$ 15 $\mu$m lying at the surface of a cell of radius R $\approx 1\,\mu$m, we estimate a bending energy per filament (see eqn. 10.5 in [40]) of

$$E_{bend} \approx \frac{L_p\, l}{2R^2} k_B T \approx 1\, k_B T \approx 4\, \mathrm{pN} \cdot \mathrm{nm}, \tag{5}$$

comparable to thermal energy and to the rough energy scale of a myosin step (a few pN·nm, estimated from a stall force of $\approx$ 0.5 pN [41] and a step size of 5 nm [37]). Given a density of 100 filaments per $\mu m^2$ of membrane, this gives us a non-negligible energy of several hundred pN·nm per $\mu m^2$. Thus, our first new term captures a driving force that rotates filaments away from the local direction of maximal curvature, favoring filament alignment in the least-curved orientation.

In the remainder of this subsection, we derive this curvature penalty term. We seek to describe the dynamics of reorientation of the filaments as they move away from the unfavored direction of maximum curvature. A phenomenological example of a curvature term that leads to reordering of the actin filaments was used in the work of Woodhouse and Goldstein on filament ordering in the giant internodal cells of algae *Chara* [42]. We consider a similar term, which describes the relaxation dynamics of filament rotation away from the direction of maximum curvature, toward the direction of minimum curvature. Conceptually, this curvature penalty term has three components,

$$\text{curvature update} = (\text{kinetic coefficient}) \cdot (\text{elastic driving force}) \cdot (\text{direction of rotation}). \tag{6}$$

The kinetic coefficient, which we will call $\varepsilon$, is a constant that tunes the contribution of the curvature term to the full Toner-Tu dynamics. The elastic driving force, which we will call $F_\kappa(\mathbf{v}, \kappa)$, computes a curvature-dependent force magnitude as a function of the orientation of vector $\mathbf{v}$ and the local curvature. Lastly, the "direction of rotation" is a geometric function that computes a vector which is orthogonal to the current filament velocity vector $\mathbf{v}$ and which, when added to $\mathbf{v}$, will rotate it away from the direction of maximum curvature $\mathbf{d_1}$ without changing the magnitude of $\mathbf{v}$. This geometric function is given by $-\left(\mathbf{d}_1 - \left(\frac{\mathbf{v}}{|\mathbf{v}|} \cdot \mathbf{d}_1\right) \frac{\mathbf{v}}{|\mathbf{v}|}\right)$, as illustrated in Figure S5. Mathematically, our curvature penalty term is therefore given by

$$\left(\frac{\partial \mathbf{v}}{\partial t}\right)_{curv} = -\, \varepsilon\, F_\kappa(\mathbf{v}, \kappa) \left(\mathbf{d}_1 - \left(\frac{\mathbf{v}}{|\mathbf{v}|} \cdot \mathbf{d}_1\right) \frac{\mathbf{v}}{|\mathbf{v}|}\right). \tag{7}$$

We now focus on $F_\kappa(\mathbf{v}, \kappa)$, the curvature-dependent elastic driving force. To define an expression for $F_\kappa$, we follow a long-standing tradition in statistical physics in which the driving force is related to the rate of change of the free energy with respect to the geometric degrees of freedom. For simplicity,

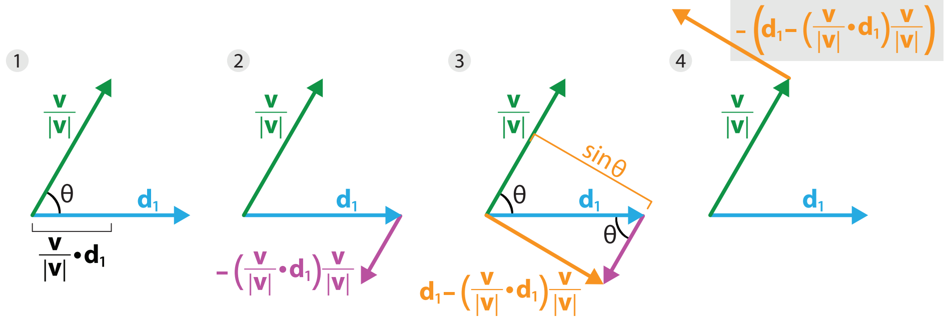

**Figure S5.** The geometric component of the curvature penalty term. This sequence provides geometric intuition for the "direction of rotation" vector which, when added to the filament velocity vector $\mathbf{v}$, will rotate it away from the direction of maximum curvature $\mathbf{d}_1$ without changing the magnitude of $\mathbf{v}$. In step 1, we note the magnitude of the dot product of two unit vectors, $(\mathbf{v}/|\mathbf{v}| \cdot \mathbf{d}_1)$. The pink vector in step 2 has this magnitude, while pointing in direction -$\mathbf{v}$. In step 3 we note that the orange vector, the sum of the blue and pink vectors, has magnitude sin $\theta$. Step 4 shows the desired "direction of rotation" vector, which acts on $\mathbf{v}$ to rotate it away from the direction of maximum curvature. We note that in the full curvature penalty term, eqn. 7, this vector is scaled by the kinetic coefficient and elastic driving force, $\varepsilon \cdot F_\kappa$.

we describe the orientation of filaments by the single parameter $\theta$, the angle measured relative to the direction of maximum curvature, $\mathbf{d}_1$. Thus, our starting point is the dynamical equation [43]

$$\frac{\partial \theta}{\partial t} = -\Gamma \frac{\partial (E_{bend})}{\partial \theta}, \tag{8}$$

a kind of relaxation dynamics which assigns a linear relationship between the temporal update of the geometric coordinate $\theta$ and the rate of change of the free energy $E_{bend}$ with respect to $\theta$ (the driving force). We can then define this energy for all orientations of an actin filament of length $L$ using an equation for elastic beam bending [44, 40],

$$E_{bend}(\theta) = \frac{EIL}{2}\kappa^2, \tag{9}$$

in which a beam of length $L$ is bent into an arc of circle of radius $R$ with curvature $\kappa = 1/R$. The combination $EI$ is known as the flexural rigidity and is the product of a material factor $E$ (the Young modulus) and a geometric factor $I$ (the areal moment of inertia).

For a general surface, the curvature tensor can be written in diagonal form as

$$\boldsymbol{\kappa} = \begin{bmatrix} \kappa_1 & 0 \\ 0 & \kappa_2 \end{bmatrix} \tag{10}$$

where $\kappa_1$ and $\kappa_2$ are the maximum and minimum curvatures, respectively, and correspond to two orthogonal directions, $\mathbf{d}_1$ and $\mathbf{d}_2$. To be clear, we can rewrite $\kappa_1 = 1/R_1$ and $\kappa_2 = 1/R_2$, where $R_1$ and $R_2$ are the radii of the circles that would best mimic our surface of interest.

To determine the bending energy as a function of filament orientation, we must know $\kappa(\theta)$, the local curvature specifically in the direction that the velocity vector is currently pointing. This quantity is a weighted average of the two principal curvatures and can be written as [45]

$$\kappa(\theta) = \kappa_1 \cos^2\theta + \kappa_2 \sin^2\theta. \tag{11}$$

Thus, the bending energy as a function of the filament orientation $\theta$ is given by

$$E_{bend}(\theta) = \frac{EIL}{2}(\kappa_1 \cos^2\theta + \kappa_2 \sin^2\theta)^2. \tag{12}$$

We now recall from eqn. 8 that the driving force we seek is linearly related to the rate of change of energy with respect to filament orientation, leading us to

$$\frac{\partial\theta}{\partial t} = -\Gamma\frac{\partial(E_{bend})}{\partial\theta} = -2\Gamma EIL\ (\kappa_1\cos^2\theta + \kappa_2\sin^2\theta)\ (\kappa_2 - \kappa_1)\cos\theta(-\sin\theta). \tag{13}$$

Folding $\Gamma$, $E$, $I$, and $L$ into the kinetic coefficient $\varepsilon$ leads us to

$$\frac{\partial\theta}{\partial t} = \varepsilon(\kappa_1\cos^2\theta + \kappa_2\sin^2\theta)(\kappa_2 - \kappa_1)\cos\theta\sin\theta, \tag{14}$$

the curvature-dependent driving force which we recall reflects a reduction in the free energy of bending due to this realignment. Finally, we combine this driving force with the direction of rotation vector, $-\left(\mathbf{d}_1 - \left(\frac{\mathbf{v}}{|\mathbf{v}|}\cdot\mathbf{d}_1\right)\frac{\mathbf{v}}{|\mathbf{v}|}\right)$. We note that, as shown in Figure S5, this vector has a magnitude of sin $\theta$. In the final combined curvature term, we want this geometric vector to be of unit length, so that the magnitude of the curvature update is set solely by the kinetic coefficient and elastic driving force. Dividing by sin $\theta$ and translating back into the language of $\mathbf{v}$, recalling that cos $\theta = \frac{\mathbf{v}}{|\mathbf{v}|}\cdot\mathbf{d}_1$, leads to

$$\left(\frac{\partial\mathbf{v}}{\partial t}\right)_{curv} = -\varepsilon\left(\kappa_1\left(\frac{\mathbf{v}}{|\mathbf{v}|}\cdot\mathbf{d}_1\right)^2 + \kappa_2\left(1 - \left(\frac{\mathbf{v}}{|\mathbf{v}|}\cdot\mathbf{d}_1\right)\right)^2\right)(\kappa_2 - \kappa_1)\left(\frac{\mathbf{v}}{|\mathbf{v}|}\cdot\mathbf{d}_1\right)\left(\mathbf{d}_1 - \left(\frac{\mathbf{v}}{|\mathbf{v}|}\cdot\mathbf{d}_1\right)\frac{\mathbf{v}}{|\mathbf{v}|}\right). \tag{15}$$

To reach the convenient shorthand used in eqn. 7, we can define

$$F_\kappa(\mathbf{v},\kappa) = \left(\kappa_1\left(\frac{\mathbf{v}}{|\mathbf{v}|}\cdot\mathbf{d}_1\right)^2 + \kappa_2\left(1 - \left(\frac{\mathbf{v}}{|\mathbf{v}|}\cdot\mathbf{d}_1\right)\right)^2\right)(\kappa_2 - \kappa_1)\left(\frac{\mathbf{v}}{|\mathbf{v}|}\cdot\mathbf{d}_1\right). \tag{16}$$

We note also that our use of the commercial finite element method software COMSOL Multiphysics® makes implementing this term relatively straightforward, as the maximum and minimal principal curvature directions and the maximum and minimum curvatures are built-in geometric variables. For the convenience of COMSOL users among our readers, we translate our variable names into the COMSOL built-ins: $\kappa_1$ = |curv1|; $\kappa_2$ = |curv2|; $\mathbf{d}_1$ = (tcurv1x, tcurv1y, tcurv1z).

The full dynamics of the velocity field in the *Toxoplasma* actin self-organization theory, now amended to account for the curvature reorientation dynamics, is given by

$$\frac{\partial\mathbf{v}}{\partial t} = [\alpha(\rho-\rho_c) - \beta|\mathbf{v}|^2]\mathbf{v} + D\nabla^2\mathbf{v} - \sigma\nabla\rho - \lambda\mathbf{v}\cdot\nabla\mathbf{v} - \varepsilon F_\kappa(\mathbf{v},\kappa)\left(\mathbf{d}_1 - \left(\frac{\mathbf{v}}{|\mathbf{v}|}\cdot\mathbf{d}_1\right)\frac{\mathbf{v}}{|\mathbf{v}|}\right). \tag{17}$$

**Toy Model of the Curvature Force: The Cylinder**

For the interested reader, in the remainder of this sub-section we consider a toy model of curvature-driven filament alignment on the surface of a simpler shape: a cylinder. This allows us to derive a specific expression for the elastic driving force in terms of the cylinder radius, $R$, rather than in the language of two arbitrary principle curvatures. On a cylindrical surface, we again characterize the orientation of filaments by the single parameter $\theta$, the angle measured relative to the circumferential direction of the cylinder. We can also define a specific curvature tensor for the cylinder,

$$\boldsymbol{\kappa} = \begin{bmatrix} 0 & 0 \\ 0 & \frac{1}{R} \end{bmatrix}. \tag{18}$$

Recalling that to work out the curvature in any direction $\mathbf{d}$ other than that of the principal curvatures, we can use the weighted average already given above as

$$\kappa = \mathbf{d}^T \boldsymbol{\kappa} \mathbf{d} = \kappa_1 \cos^2\theta + \kappa_2 \sin^2\theta. \tag{19}$$

Using this averaging operation, we find the curvature when the actin filaments are oriented at an angle $\theta$ with respect to the circumferential direction as

$$\kappa(\theta) = \frac{1}{R}\cos^2\theta. \tag{20}$$

We see that this has the right limits in that for $\theta = 0$ we recover our usual notion of the radius of the cylinder and for $\theta = \pi/2$, we find that the curvature is zero, corresponding effectively to a radius of curvature that goes to infinity.

We can now find the energy for all orientations of the actin filaments of length $L$, Young modulus $E$, and areal moment of inertia $I$, oriented at an angle $\theta$, using

$$E_{bend}(\theta) = \frac{EIL}{2}\kappa^2 = \frac{EIL}{2R^2}\cos^4\theta. \tag{21}$$

Given this energy as a function of the geometric coordinate $\theta$, we can now compute the change in free energy during reorientation of the filaments as

$$\frac{\partial(E_{bend})}{\partial\theta} = \frac{EI}{2R^2}4L\cos^3\theta(-\sin\theta). \tag{22}$$

Finally, given this expression for the energy, we can compute the dynamical equation for the evolution of $\theta$ using eqn. 8, resulting in

$$\frac{\partial\theta}{\partial t} = \Gamma\frac{2EIL}{R^2}\cos^3\theta\sin\theta. \tag{23}$$

To get a feeling for this, Figure S6 shows how the driving force depends upon the angle $\theta$. The blue curve ($\kappa_1 = 1\ \mu m^{-1}; \kappa_2 = 0\ \mu m^{-1}$) corresponds to the driving force for a cylinder. The other curves illustrate how the driving force's dependence on $\theta$ changes for different geometries, where the maximum and minimum curvatures are not as different.

*6.4. Extending flocking theory to capture Toxoplasma actin biology: filament polymerization and depolymerization*

Our second addition to the minimal Toner-Tu theory are density source and sink terms, which are usually not included for macroscopic flocks or herds. Although the total number of animals in a school, flock, or herd can of course change over time as animals are born and die, the time scales over which these changes happen are long compared to the time scale of the flocking behavior itself. They can thus be neglected when accounting for macroscopic changes to density or velocity fields during flocking. In the case of *Toxoplasma* actin filament self-organization, however, both the time scales of actin filament polymerization and depolymerization (see Appendix Section 10.2) and the time scale of filament transport across the entire cell surface are on the order of seconds (see Section 7.2). Moreover, we know that *T. gondii* actin polymerization is favored specifically at the anterior end of the cell (the conoid) by the formin FRM1 [46, 18], and that filaments are further stabilized by the actin-binding protein GAC during anterior microneme secretion events at the anterior end [18]. Conversely, *T. gondii* actin depolymerization is promoted by proteins like profilin [47] and actin depolymerizing factor (ADF) [10], which are not known to be spatially restricted.

In order to mathematicize this biological intuition and capture the short time scale dynamics of actin filaments, we amend the continuity equation. In its simplest form, the continuity equation acknowledges only one way to change the quantity of molecules in a material volume element: by molecules moving in

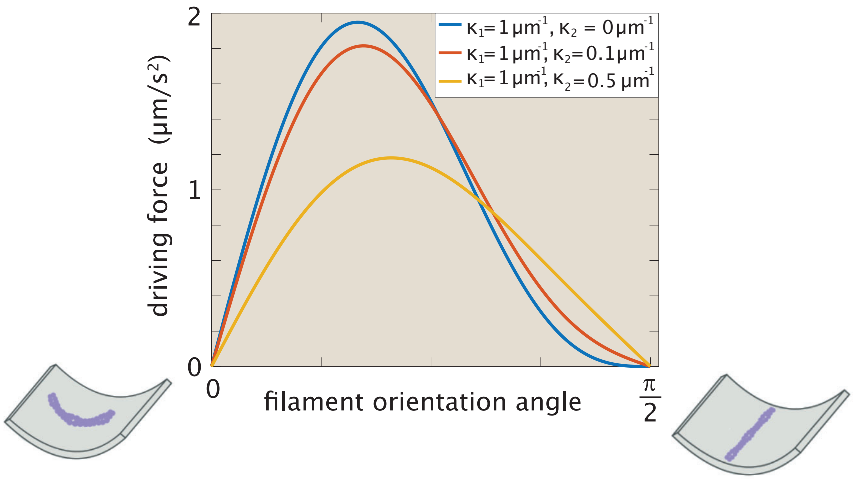

**Figure S6.** Physics of reorientation of actin filaments due to curvature. The graph shows the driving force for reorientation ($\partial\theta/\partial t$, eqn. 14, with $\varepsilon = 6\ \mu m^3/s^2$) as a function of the filament orientation angle $\theta$ with respect to the direction of maximum curvature. The blue curve shows the driving force anywhere on a cylindrical surface, where $\kappa_2 = 0$, and $\theta$ is defined with respect to the circumferential direction. The red and yellow curves show the driving force at a point on any arbitrary surface with local curvature defined by $\kappa_1$ and $\kappa_2$.

and out of that element, to and from neighboring volume elements. To incorporate the reality of filament polymerization and depolymerization, we must allow sources and sinks of filament density. At the cell anterior, we add both a source term $c$ and a degradation term $-\gamma\rho$, leading to an amended version of the continuity equation that can be written as

$$\frac{\partial\rho}{\partial t} = -\nabla\cdot(\rho\mathbf{v}) - \gamma\rho + c. \tag{24}$$

Actin filament density is added (polymerized) at a constant rate $c$ and lost (depolymerized) at a density-dependent rate $\gamma\rho$. Over the rest of the cell surface, outside the conoid, polymerization is not favored but depolymerization still occurs. Thus,

$$\frac{\partial\rho}{\partial t} = -\nabla\cdot(\rho\mathbf{v}) - \gamma\rho. \tag{25}$$

The Toner-Tu equations in conjunction with these generalized versions of the continuity equation constitute the full conceptual formulation of the problem of actin dynamics for *Toxoplasma*, as summarized in Figure S3.

#### 7. Parameter Choices and Dimensionless Ratios

One tool for developing intuition for partial differential equations, such as the *Toxoplasma* actin self-organization equations presented above, is to reformulate these governing equations in dimensionless form. In this section, we first explore the dimensionless version of our equations and develop intuition for the dimensionless ratios they present. We then estimate and define parameter values for the specific case of the *Toxoplasma gondii* actin surface layer in “real-world” units. A parallel discussion of the dimensionless Toner-Tu equations and parameter estimates for macroscopic herding animals is presented in our manuscript on wildebeest herding on arbitrary curved surfaces [8].

*7.1. Dimensionless ratios in the theory*

We seek here to explain in an accessible way the process of reformulating the governing equations in dimensionless form. We begin by explicitly defining the units of the two key field variables $\rho$ and $\mathbf{v}$. The density field describes the number of filaments found per unit area in the thin surface layer of the *Toxoplasma* cells. This implies that we have

$$[\rho] = \frac{1}{L^2}, \tag{26}$$

where we use the notation that [*thing*] means "units of thing," and $L$ and $T$ are units of length and time, respectively. Similarly, the field $\mathbf{v}$ has units given by

$$[v] = \frac{L}{T}. \tag{27}$$

With these definitions in hand, we can now explore the units of the various terms that appear in the Toner-Tu equations given in eqn. 17, and then define dimensionless variables that will allow us to recast the equations in dimensionless form. The time derivative of the $\mathbf{v}$-field results in a quantity with units

$$[\frac{\partial v_i}{\partial t}] = \frac{L}{T^2}. \tag{28}$$

Thus, all the other terms in the Toner-Tu equations must have these same units. We can determine the units of the advection term using

$$[\lambda v_j \frac{\partial v_i}{\partial x_j}] = \frac{L}{T^2}. \tag{29}$$

For this to be true, we must have

$$\lambda \times \frac{L}{T} \times \frac{1}{T} = \frac{L}{T^2} \tag{30}$$

which implies that $\lambda$ is dimensionless,

$$[\lambda] = 1. \tag{31}$$

To evaluate the units in the velocity selection term we note that we have

$$[\alpha \times \rho \times v] = [\alpha] \times \frac{1}{L^2} \times \frac{L}{T} = \frac{L}{T^2} \tag{32}$$

which implies that

$$[\alpha] = \frac{L^2}{T}. \tag{33}$$

Similar reasoning allows us to determine the units of the quantity $\beta$ using

$$[\beta v^3] = \beta \frac{L^3}{T^3} = \frac{L}{T^2}. \tag{34}$$

This implies that

$$[\beta] = \frac{T}{L^2}. \tag{35}$$

The neighbor coupling diffusion-like term tells us

$$D\frac{\partial^2 v}{\partial x^2} = D\frac{L/T}{L^2} = \frac{L}{T^2} \tag{36}$$

which gives us the dimensions of $D$ as

$$D = \frac{L^2}{T}. \tag{37}$$

We next tackle the pressure term which requires

$$[\sigma \frac{\partial \rho}{\partial x}] = \sigma \frac{1}{L^3} = \frac{L}{T^2}. \tag{38}$$

This implies that

$$[\sigma] = \frac{L^4}{T^2}. \tag{39}$$

Lastly, the curvature term demands that

$$\varepsilon \frac{1}{L^2} = \frac{L}{T^2} \tag{40}$$

which lead us to conclude that

$$[\varepsilon] = \frac{L^3}{T^2}. \tag{41}$$

We now have a clear picture of the units of each of the material parameters that appears in the Toner-Tu *T. gondii* actin self-organization theory, and we move on to defining our dimensionless variables. For example, for the positional coordinate, we take

$$x^* = \frac{x}{L}, \tag{42}$$

where $L$ is a characteristic length scale in the problem, which we take to be the cell size, $L \approx 5$ $\mu$m. Similarly, we use a characteristic velocity scale $U \approx 5$ $\mu$m/s, set by myosin A speed, to define the dimensionless velocity as

$$v^* = \frac{v}{U}. \tag{43}$$

The density can be rescaled by using the critical density $\rho_c$ as our scaling variable, resulting in

$$\rho^* = \frac{\rho}{\rho_c}. \tag{44}$$

Recalling that curvature has units of $1/L$, we define our scaled curvature variable through

$$\kappa^* = L\kappa. \tag{45}$$

Lastly, in light of the definitions above, we can define a time scale $L/U$, the time it takes for a myosin-transported actin filaments to cross the entire cell, culminating in the definition

$$t^* = \frac{t}{L/U}. \tag{46}$$

Using the definitions given above, we can now rewrite the Toner-Tu actin self-organization equations using the dimensionless versions of $t$, $x$, $\rho$ and $\mathbf{v}$ as

$$\frac{U^2}{L}\frac{\partial v_i^*}{\partial t^*} = \rho_c U \alpha(\rho^* - 1)v_i^* - \beta U^3 |\mathbf{v}^*|^2 v_i^* - \frac{\sigma \rho_c}{L}\frac{\partial \rho^*}{\partial x_i^*} + \frac{DU}{L^2}\nabla_*^2 v_i^* - \frac{\lambda U^2}{L} v_j^* \frac{\partial v_i^*}{\partial x_j^*} - \frac{\varepsilon}{L^2} F_\kappa(v^*, \kappa^*)\left((d_1)_i - \frac{v_i^* v_j^* (d_1)_j}{v_k^* v_k^*}\right). \tag{47}$$

We then divide everything by $U^2/L$, resulting in five dimensionless parameters and full dynamical equations of the form

$$\frac{\partial v_i^*}{\partial t^*} = \frac{\rho_c L \alpha}{U}(\rho^* - 1)v_i^* - LU\beta |\mathbf{v}^*|^2 v_i^* - \frac{\sigma \rho_c}{U^2}\frac{\partial \rho^*}{\partial x_i^*} + \frac{D}{UL}\nabla_*^2 v_i^* - \lambda v_j^* \frac{\partial v_i^*}{\partial x_j^*} - \frac{\varepsilon}{LU^2} F_\kappa(v^*, \kappa^*)\left((d_1)_i - \frac{v_i^* v_j^* (d_1)_j}{v_k^* v_k^*}\right). \tag{48}$$

To write the dimensionless version of the continuity equation, we use the same definitions of dimensionless variables given above and the same strategy for replacing dimensionful variables with their dimensionless counterparts. We find that the generalized continuity equation takes the form

$$\frac{\partial \rho^*}{\partial t^*} = -\nabla_* \cdot (\rho^* v^*) - \frac{\gamma L}{U}\rho^* + \frac{L}{U\rho_c}c. \tag{49}$$

Writing the Toner-Tu equations in this form gives us a handle on the meaning of the different terms. The two components of the velocity selection term have dimensionless parameters given by

$$\text{velocity selection } \alpha \text{ term} = \frac{\alpha \rho_c L}{U} = \frac{L/U}{1/(\alpha\rho_c)} = \frac{\text{time for filament to cross cell}}{\text{time for speed to increase to } v_{myosin}} \tag{50}$$

and

$$\text{velocity selection } \beta \text{ term} = \beta L U = \frac{L/U}{1/(\beta U^2)} = \frac{\text{time for filament to cross cell}}{\text{time for speed to decrease to } v_{myosin}}. \tag{51}$$

Each of these terms provides intuition about how quickly the filaments return to the steady state mean-field speed $v_{myosin}$ given some perturbation that disturbs them from that value. Recall that $L/U$ (the "time for filament to cross cell") is roughly the time it takes for an actin filament to be transported by myosin across the length of the cell. The pressure term can be rewritten as

$$\text{pressure term} = \frac{\rho_c}{U^2/\sigma} = \frac{\text{critical density}}{\text{typical density excursion away from } \rho_0}. \tag{52}$$

Increasing $\sigma$ decreases density variance; in other words, densities that emerge are within a more narrow range around the mean density $\rho_0$. The neighbor coupling term that carries out democratic velocity smoothing has the dimensionless prefactor

$$\text{democracy term} = \frac{D}{UL} = \frac{L/U}{L^2/D} = \frac{\text{time for filament to cross cell}}{\text{time for velocity to diffuse across cell}}, \tag{53}$$

analogous to the Péclet number. The velocity self-advection term prefactor $\lambda$, which for our case is 1, can be written as

$$\text{advection term} = \frac{\lambda L/U}{L/U} = \frac{\text{time for velocity advection across cell}}{\text{time for filament to cross cell (density advection)}}. \tag{54}$$

Finally, the curvature penalty term can be written as

$$\text{curvature penalty ratio} = \frac{\varepsilon}{LU^2} = \frac{L/U}{UL^2/\varepsilon} = \frac{\text{time for filament to cross cell}}{\text{time for filament to rotate away from maximum curvature}}. \tag{55}$$

To interpret this ratio, we remember that $\kappa = 1/L$ is a typical cell curvature, that the curvature-induced elastic driving force scales with $\kappa^2$, and that $\mathbf{v}/U$ is a unit vector indicating the *orientation* of a filament. Thus, $\varepsilon/UL^2$ gives a curvature-dependent rate of rotation or relaxation for filament orientation, and $UL^2/\varepsilon$ gives a time scale for curvature-dependent rotation.

For each of these cases, the value of the dimensionless ratio gives us a sense of how large a contribution the term of interest will make in the incremental update to $\mathbf{v}(t)$. That is, if we think of the numerical solution to the equations, during every time step $\Delta t$, $\mathbf{v}(t)$ will get updated at every point in space. How much $\mathbf{v}(t)$ changes depends upon all the different contributions in the governing equation. These dimensionless ratios measure the relative importance of each term, serving roles analogous to the Reynolds number in thinking about the Navier-Stokes equations and the Péclet number in the context of coupled diffusion-advection problems.

Next, we turn to the two dimensionless ratios in the density governing equation. The term multiplying $\rho^*$ on the right side of eqn. 49 involves the ratio of two very important time scales, namely,

$$\frac{\text{time for filament to cross cell}}{\text{time to depolymerize actin filament}} = \frac{L/U}{1/\gamma}. \tag{56}$$

Intuitively, we expect that when this dimensionless ratio is of order unity or larger ($\gamma > U/L$), then a rearward steady-state flow can be achieved because the actin filaments do not live long enough to accumulate at the cell posterior and force a return orbit.

The actin production rate is also given by a dimensionless ratio of the form

$$\frac{\text{actin density production rate}}{\text{actin density transport rate}} = \frac{c}{\rho_c U/L}. \tag{57}$$

*7.2. Parameter choices for modeling Toxoplasma actin self-organization*

In this subsection, we estimate parameter values for the specific case of *Toxoplasma gondii* surface F-actin organization, and we define the parameter values used in the simulations reported in the main text. We also discuss the transition from the recirculating to the unidirectional F-actin state, using estimates and calculations to sanity check our parameter choices for filament polymerization and depolymerization. To determine our parameters, we use our knowledge of *Toxoplasma gondii* actin biology, order-of-magnitude estimates, and a principle of balancing terms.

Based on published electron microscopy of actin filaments from gliding *Toxoplasma gondii* [9], we estimate a rough density of 150 filaments/$\mu m^2$. This average density will serve as the initial uniform density in our simulations,

$$\rho_0 = 150\frac{1}{\mu m^2}. \tag{58}$$

We next estimate $\rho_c$, the critical density above which actin filaments interact with their neighbors and collectively organize. Actin filaments in *T. gondii* are thought to be of order 100 nm in length [48, 49]. To estimate the critical density, we consider the density scale at which filaments can explicitly interact with neighboring filaments. If we imagine that each filament is free to rotate, sweeping out a circular area with diameter 100 nm, these circles will begin to overlap when more than 10 x 10 filaments are packed within a 1 $\mu$m x 1 $\mu$m square. Thus, we estimate that filaments collectively organize above a density

$$\rho_c = 100\frac{1}{\mu m^2}. \tag{59}$$

To get a feel for this density, picture 10 rows of 10 filaments, each of length 0.1 $\mu$m, in a 1 $\mu$m by 1 $\mu$m square.

We next move to the parameters of the governing equation for velocity, where a principle of balancing terms will play an important role. That is, in order to explore the contributions of all terms in the velocity equations, we balance their magnitudes to ensure that each contributes roughly equally to the overall velocity update. To get an initial estimate of the scale of that update, we estimate the typical change in velocity as

$$\frac{\partial v}{\partial t} \approx \frac{10\ \mu m/s}{1\ s} \approx 10\ \mu m/s^2, \tag{60}$$

considering that a filament reaching the end of the cell at roughly 5 $\mu m/s$ will reverse directions (and head back up the cell at velocity -5 $\mu m/s$) over a time scale of roughly one second. Hence, the acceleration is estimated to be $\approx 10\ \mu m/s^2$. Now that we have this as the typical scale of the change in velocity, we can examine each term in the governing equation and determine its material parameter by demanding that it yield a contribution of comparable magnitude. Let's try this out for the velocity selection terms. First, we have

$$10\ \mu m/s^2 = \alpha(\rho - \rho_c)v \approx \alpha \times 50\ \frac{1}{\mu m^2} \times 5\ \mu m/s \tag{61}$$

which leads to the conclusion that

$$\alpha = 0.04\ \mu m^2/s. \tag{62}$$

To determine the parameter $\beta$ we adopt the same strategy, with

$$10\ \mu m/s^2 = \beta v^3 \approx \beta \times (5\ \mu m/s)^3 \tag{63}$$

which leads to the conclusion that

$$\beta = 0.08\ s/\mu m^2. \tag{64}$$

We can now apply this thinking to make an estimate of the neighbor coupling coefficient using the equality

$$10\ \mu m/s^2 = D\frac{\partial^2 v}{\partial x^2} \approx D\frac{5\ \mu m/s - (-5\ \mu m/s)}{(1\ \mu m)^2}, \tag{65}$$

where we again imagine a change in filament velocity from 5 $\mu m/s$ to -5 $\mu m/s$ and take 1 $\mu$m to be the length scale of such gradients in the velocity, leading to an estimate for the neighbor coupling coefficient of

$$D = 1\ \mu m^2/s. \tag{66}$$

To find the parameter $\sigma$ that tunes the pressure term, we estimate a density gradient scale of $\rho - \rho_c = 50\ 1/\mu m^2$ and make the correspondence

$$10\ \mu m/s^2 = \sigma\frac{\partial \rho}{\partial x} \approx \sigma\frac{50\ \frac{1}{\mu m^2}}{1\ \mu m} \tag{67}$$

which leads to a choice of

$$\sigma = 0.2\ \mu m^4/s^2. \tag{68}$$

Our final term in the Toner-Tu equations themselves concerns the penalty for filament alignment in a direction of high curvature. Recalling eqn. 15 and estimating values of 4 $\mu m^{-1}$ for $\kappa_1$, 1 $\mu m^{-1}$ for $\kappa_2$, and $1/\sqrt{(}2)$ for $\frac{\mathbf{v}}{|\mathbf{v}|} \cdot \mathbf{d}_1$, we have

$$10\ \mu m/s^2 \approx -\varepsilon(2.5\frac{1}{\mu m})(1\frac{1}{\mu m} - 4\frac{1}{\mu m})(\frac{1}{\sqrt{2}})(1 - \frac{1}{\sqrt{2}}) \tag{69}$$

which leads us to

$$\varepsilon = 6\ \mu m^3/s^2. \tag{70}$$

The advection term warrants special discussion. Following the tradition of Toner and Tu [34], the dimensionless coefficient $\lambda$ does not need to be equal to 1 for the case of flocking animals. If $\lambda = 1$, this term simply enforces the material time derivative; in other words, the velocity field advects itself, as in the Navier-Stokes equation. Flocking birds or sheep, however, can display behaviors - like slowing down when they notice a gradient of decreasing velocity in the birds ahead of them - that modify the parameter $\lambda$ away from a value of 1. In order to show the theory in its most generally useful form, we have maintained $\lambda$ throughout as a tunable parameter. However, for our simulations of actin filament self-organization, we do not hypothesize any behavioral response to the velocity gradient, as described above for bird flocks. Thus, we have

$$\lambda = 1. \tag{71}$$

Using a principle of balancing terms, we have made a first guess at each of the parameters appearing in the Toner-Tu formulation for *Toxoplasma.* Note that the main purpose of the use of the Toner-Tu theory in the context of our work on *Toxoplasma* was to understand what kinds of behaviors are possible. From that point of view, getting a sense of the characteristic scales of the different parameters suffices. That said, we can imagine explicit experimental strategies aimed at measuring and ultimately controlling the parameters in the theory, and we believe this represents an exciting direction for future work.

Finally, we turn to estimating the parameters $c$ and $\gamma$ of the equation for actin filament density,

$$\frac{\partial \rho}{\partial t} = -\nabla \cdot (\rho \mathbf{v}) - \gamma \rho + c, \tag{72}$$

where the two terms on the far right govern depolymerization and polymerization of filaments, respectively. From the previous section, we remember a key dimensionless ratio,

$$\frac{\text{time for filament to cross cell}}{\text{time to depolymerize actin filament}} = \frac{L/U}{1/\gamma}. \tag{73}$$

Intuitively, we expect that a transition between recirculating and unidirectional F-actin flow could arise when this dimensionless ratio is of order unity. Thus, we can estimate that

$$\gamma = \frac{U}{L} \approx \frac{5\ \mu\text{m/s}}{5\ \mu\text{m}} \simeq 1\ s^{-1}, \tag{74}$$

or a bit smaller, given our rounding down of cell length. In Figure 4, we find interesting changes in behavior as we sweep through a range of values for $\gamma$ from 0.2 $s^{-1}$ to 0.7 $s^{-1}$, and in our estimate for $c$ below, we use a value of $\gamma \approx 0.5\ s^{-1}$.

The actin filament polymerization rate is also given by a dimensionless ratio of the form

$$\frac{\text{actin density production rate}}{\text{actin density transport rate}} = \frac{c}{\rho_c U/L}. \tag{75}$$

A rough estimate of the parameter $c$ can be gotten by considering a simplified scenario in which density is uniform across the cell surface, and is in steady state, unchanging in time. Importantly, we must remember that actin polymerization is promoted specifically at the cell anterior, in the *T. gondii* conoid [46, 18]. In our model, filament density production at rate $c$ is restricted to the conoid region, as shown in Supplementary Figure S3. This region has a surface area $A_{conoid}$, which is approximately 1/20th of $A_{tot}$, the total surface area of the cell. Thus, the steady state density balance for filaments will take the form

$$-\gamma \rho_{ss} A_{tot} + c A_{conoid} = 0. \tag{76}$$

We can solve this for the steady-state filament density,

$$\rho_{ss} = \frac{c}{\gamma} \frac{A_{conoid}}{A_{tot}}. \tag{77}$$

Setting the steady-state filament density equal to our estimated average density, $\rho_0$, we obtain an estimate for our filament polymerization coefficient,

$$c = \gamma \rho_0 \frac{A_{tot}}{A_{conoid}} \approx 0.5\ s^-1 \times 150\ \mu m^-2 \times 20 \approx 1500\ \mu m^{-2}\ s^{-1}. \tag{78}$$

We use this value for $c$ in Figure 4A, and in Figure 4B we sweep through a range of values for $c$ from 0 to 2000 $\mu m^{-2}\ s^{-1}$. In Section 10.2 of the Appendix, we compare the rates of filament polymerization and depolymerization determined above to data and estimates on actin dynamics in well-studied biological systems, and we find them to be similar. While the rates proposed above seemed large to us at a first glance, the more considered deliberation presented in the Appendix convinced us that these rates are in a reasonable range.

#### 8. Deriving a Tangential Formulation of the Filament Self-Organization Equations for a Curved Surface

Thus far, our treatment of the actin filament self-organization equations presented here has ignored the fact that in our *T. gondii* case of interest, the "flock" of actin filaments is moving around on the curved surface of the cell. Our mathematics must account for this geometry; for example, the evaluation of the derivatives used in the governing partial differential equations requires a knowledge of the local curvature. In this section, we recast our governing equations in a tangential formulation for curved surfaces, using an extrinsic differential geometry approach rather than a more traditional intrinsic one. The presentation here is abridged; for a more thorough discussion of this general curved-surface reformulation of the Toner-Tu theory, we refer the reader to our manuscript on animal herding on arbitrary curved surfaces [8].

Traditional formulations of the differential geometry of surfaces begin with the parametrization of a surface of interest using a series of points given by $\mathbf{r}(u, v)$, where $u$ and $v$ are the parameters that characterize the surface of interest. The surface of a cylinder of radius $R$, for example, can be represented by

$$\begin{aligned}\mathbf{r}(\theta, z) &= (x(\theta, z), y(\theta, z), z(\theta, z)) \\ &= (R\cos\theta, R\sin\theta, z).\end{aligned} \tag{79}$$

However, the complex shape of the *Toxoplasma gondii* cells of interest here does not conform to the simple parameterization of highly idealized geometries. While motile *Toxoplasma gondii* tachyzoite shape is stereotypical and consistent across cells, it is highly asymmetric, with features like a crescent-like bend and a narrow protrusion (the conoid) at the anterior cell end. In order to understand the contributions of the real-world, biological geometry of these cells, we want to solve our self-organization equations on their true asymmetric shapes. This is clearly not possible analytically.

To move toward an ultimate goal of solving our equations numerically using the finite element method, we can reformulate them using an extrinsic differential geometry approach; we carry out the mathematics in the full three-dimensional setting of $\mathbb{R}^3$, while using a knowledge of the normal vector $\mathbf{n} = (n_1, n_2, n_3)$ everywhere on the surface of interest to project the governing equations onto the surface. For a full description of the tangential surface formulation of flocking governing equations on an arbitrary curved surface, the reader is directed to our recent preprint [8]. Central to this extrinsic geometry approach is the projection operator, defined as

$$P_{ij} = \delta_{ij} - n_i n_j. \tag{80}$$

The curved space or tangential version of the pressure term $\sigma\nabla\rho$, for example, requires projecting the full 3D gradient of the density onto the tangent plane. We follow Jankuhn *et al.* [50] in introducing the notation $\nabla_\Gamma$ for the projection of the gradient onto the surface of interest. Mathematically, this amounts to computing

$$(\nabla_\Gamma \rho)_i = (\delta_{ij} - n_i n_j)\,\frac{\partial \rho}{\partial x_j}. \tag{81}$$

We again refer the reader to [8] for a detailed explication of the use of tangential differential operators to generate surface-projected versions of all the other terms in the Toner-Tu equations. Here, we modify the curvature penalty term to its curved-surface implementation, since it is an original contribution to the *Toxoplasma* actin self-organization equations and not included in reference [8]. We first define $\mathbf{v}^{\|}$, the in-plane velocity,

$$v_i^{\|} = P_{ij} v_i = v_i - n_i v_i n_j. \tag{82}$$

The curved-surface formulation of the curvature penalty update, derived for the plane in Section 6.3, can now be written in component form as

$$\left(\frac{\partial v_i^{\|}}{\partial t}\right)_{\Gamma,curv} = -\varepsilon\, F_\kappa(v_i^{\|},\kappa)\left((d_1)_i - \frac{v_i^{\|}v_j^{\|}(d_1)_j}{v_k^{\|}v_k^{\|}}\right). \tag{83}$$

The tangential curved-surface form of the full governing equations for velocity is

$$\begin{aligned}\frac{\partial v_i^{\|}}{\partial t} &= [\alpha(\rho-\rho_c) - \beta v_j^{\|}v_j^{\|}]v_i^{\|} - \sigma\left(\frac{\partial \rho}{\partial x_i} - n_i n_j \frac{\partial \rho}{\partial x_j}\right) + DP_{il}\left(\frac{\partial G_{lj}}{\partial x_j} - n_j n_k \frac{\partial G_{lj}}{\partial x_k}\right) \\ &- \lambda\left[\left(\frac{\partial v_i^{\|}}{\partial x_j} - n_l n_j \frac{\partial v_i^{\|}}{\partial x_l}\right) - n_i n_k\left(\frac{\partial v_k^{\|}}{\partial x_j} - n_l n_j \frac{\partial v_k^{\|}}{\partial x_l}\right)\right] v_j^{\|} - \varepsilon\, F_\kappa(v_i^{\|},\kappa)\left((d_1)_i - \frac{v_i^{\|}v_j^{\|}(d_1)_j}{v_k^{\|}v_k^{\|}}\right),\end{aligned} \tag{84}$$

where $\mathbf{d}_1$ is a unit vector in the direction of maximum local curvature, $F_\kappa$ is defined in eqn. 16, and $\mathbf{G}$ is the surface velocity gradient tensor which in component form can be written as

$$G_{ij} = \left(\frac{\partial v_i^{\|}}{\partial x_j} - n_l n_j \frac{\partial v_i^{\|}}{\partial x_l}\right) - n_i n_k\left(\frac{\partial v_k^{\|}}{\partial x_j} - n_l n_j \frac{\partial v_k^{\|}}{\partial x_l}\right). \tag{85}$$

Similarly, the curved-surface formulation of the governing equation for density can be written as

$$\frac{\partial \rho}{\partial t} = -\frac{\partial(\rho v_i^{\|})}{\partial x_i} + n_i n_j \frac{\partial(\rho v_i^{\|})}{\partial x_j} - \gamma\rho + c. \tag{86}$$

We now have a complete formulation of our self-organization governing equations for the general surface context, requiring only a description of the surface in the language of normal vectors.

#### 9. Numerically Solving the Filament Self-Organization Equations on the *Toxoplasma* Cell Surface

In order to solve the curved-surface formulation of our actin filament self-organization equations for general initial conditions and on general geometries, like the surface of the *Toxoplasma gondii* cell, we turn to the finite element method (FEM). More specifically, we make use of the custom PDE solver capabilities of the commercial FEM software COMSOL Multiphysics® [51]. The finite element method enables us to represent the cell surface as a mesh of triangles and solve our dynamical equations at the nodes of those triangles, then interpolate between them. In this section, we first detail the creation of surface meshes compatible with the finite element method from the soft X-ray tomograms of extracellular *Toxoplasma* tachyzoites described in Section 4.7. We then discuss our use of the finite element method to numerically solve the tangential self-organization equations on these surface meshes.

##### *9.1. Spherical harmonic shape analysis of Toxoplasma cells and mesh generation*

Many fields of study are now dependent upon fast and reliable representations of surfaces. One particular tool for doing so, called SPHARM-PDM, converts image segmentations into a mathematical representation of shape in the language of spherical harmonics, and then into triangulated surfaces [52]. This approach has proven useful for shape analysis and comparison in the macroscopic contexts of archaeology and medical imaging [53], and in this study we use it at microscopic scales, to describe the shape of single cells. In brief, the SPHARM tool maps the closed surface of an object of interest (which must have spherical topology) to the surface of a sphere, preserving area and minimizing distortion [52]. This mapping enables a mathematical description of the surface of interest as a series of spherical harmonics [54]. In essence, the SPHARM description is a list of coefficients, which weight a series of basis

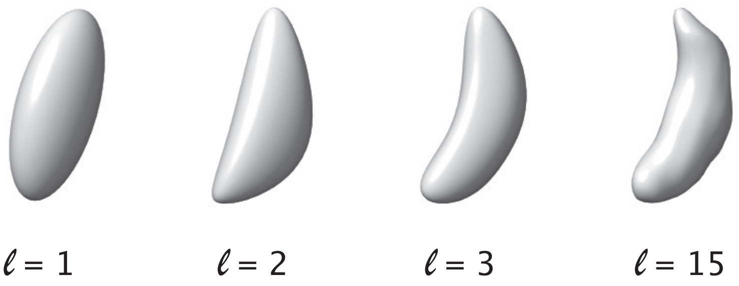

**Figure S7.** The surface of an extracellular *T. gondii* tachyzoite cell, represented by a series of spherical harmonic modes. Each panel shows the cell shape representation when carried to the order of expansion indicated by the value of $\ell$. For example, the right-most representation is $\mathbf{r}(\theta,\varphi)=\sum_{l=0}^{15}\ \sum_{m=-l}^{l}\ c_{lm}^{(i)}\ \ Y_{lm}(\theta,\varphi)$.

functions: the spherical harmonics. The spherical harmonics $Y_{lm}(\theta,\varphi)$ are solutions to Laplace's equation on the surface of a sphere and form a "basis" from which we can represent any function parametrized by $\theta$ and $\phi$ as

$$\mathbf{r}(\theta,\varphi)=\begin{bmatrix} x(\theta,\varphi)\\ y(\theta,\varphi)\\ z(\theta,\varphi)\end{bmatrix}=\sum_{l=0}^{\infty}\sum_{m=-l}^{l}\begin{bmatrix} c_{lm}^{(x)}\\ c_{lm}^{(y)}\\ c_{lm}^{(z)}\end{bmatrix}\quad Y_{lm}(\theta,\varphi). \tag{87}$$

All the character of a given closed surface is captured in the coefficients $c_{lm}^{(i)}$, where the superscript $i$ tells us which component of the vector $\mathbf{r}(\varphi,\theta)$ is being considered. In other words, different closed surfaces are described using the same series of harmonics $(Y_{lm}(\theta,\varphi))$, but the series of coefficients $c_{lm}^{(i)}$ will differ, giving more or less weight to a given harmonic in the overall representation of a given surface. To obtain these coefficients, the orthonormality of the spherical harmonics is used to pick off a given coefficient by multiplying both sides of eqn. 87 by $Y_{l'm'}^{*}(\theta,\varphi)$ and integrating over $\theta$ and $\varphi$ to obtain

$$c_{\mathrm{l'm'}}^{(x)}=\int_0^{2\pi}d\varphi\int_0^{\pi}\sin\theta\ d\theta\quad Y_{l'm'}^{*}(\theta,\varphi)x(\theta,\varphi). \tag{88}$$

Using these ideas, the surface of a representative *Toxoplasma gondii* tachyzoite cell was broken up into a series of weighted spherical harmonic modes. We note that *Toxoplasma gondii* tachyzoites have a consistent, stereotyped shape and size. First, reconstructed 3-dimensional soft X-ray tomography images of extracellular, activated *T. gondii* tachyzoite cells were segmented in ChimeraX [55] and converted into binary image stacks. Within the 3D Slicer package[56], images of one representative cell were converted to a label map read in by the SPHARM-PDM Generator module, used within the SlicerSalt package [57]. As described above, a SPHARM shape description was generated in the language of spherical harmonics. Including a larger number of modes in the shape description will better capture the original surface shape, as shown in Figure S7. For our representations, we used $\ell = 15$ modes. The surface thus represented was sampled into a triangular mesh (PDM), which is compatible with the finite element method and was used to solve the governing partial differential equations for the density and velocity fields within COMSOL Multiphysics®.

##### *9.2. Solving the tangential self-organization equations in COMSOL Multiphysics®*

To solve our custom surface partial differential equations (eqns. 84 and 86), we used the COMSOL Multiphysics® General Form Boundary PDE interface and took advantage of COMSOL's built-in tangential differentiation operator, dtang(f,x). We also made use of the normal vector (nx, ny, nz), a built-in geometric variable, and the curvature variables discussed in Section 6.3. For a thorough and practical introduction to the finite element method, we recommend reference [58]. For additional details on rewriting the curved-surface Toner-Tu equations in a form convenient for standard finite element method solvers and for implementing specifically in COMSOL Multiphysics®, we refer the reader to reference [8]. For each simulation, we initialized with uniform density field $\rho_0 = 150\ 1/\mu m^2$ and a disorganized velocity field, in which every node's velocity orientation was drawn randomly from a uniform distribution of angles between 0 and $2\pi$ and had magnitude $v(0) = 5\ \mu\text{m/s} \approx v_{myosin}$. Other parameters are defined and discussed in detail in Section 7. Our standard mesh size involved 660-794 triangular elements. To avoid the accumulation of out-of-plane components in $\mathbf{v}$ from numerical error, we implemented a weak constraint of $\mathbf{n} \cdot \mathbf{v} = 0$. Similarly, in the model with no filament turnover, we implemented a global constraint on the total integrated density $\rho$ on the surface, ensuring it stayed at its initial value. Default COMSOL solvers and settings were used: implicit backward differentiation formula (BDF) for time stepping and multifrontal massively parallel sparse direct solver (MUMPS) for the linear direct spatial solver.

#### 10. Appendix

##### *10.1. Dry vs. wet active matter: estimating frictional drag from a fixed surface*

As discussed in section 6.1, the field of active matter has distinguished so-called wet and dry active matter. Qualitatively, the distinction centers on whether there is hydrodynamic coupling between the active agents through the intervening medium that they occupy. In wet active matter, fluid mediated coupling between active agents is significant. In dry active matter, these viscous forces are negligible relative to frictional drag, commonly due to the presence of a "no slip" boundary, as when filaments move over a fixed glass slide during a traditional *in vitro* gliding assay. To help assess whether the dynamics of our *Toxoplasma gondii* surface actin are in the wet or dry regime, we sought an order-of-magnitude estimate of frictional drag, for comparison to viscous forces.

For an actin filament moving in the thin $\approx 25$ nm layer between the plasma membrane and the inner membrane complex, we can compute the way that the filament entrains the surrounding fluid using the driven Stokes equations with friction,

$$-\eta\nabla^2\mathbf{v} + \nabla\pi = \mathbf{F}_{myosin} - \gamma\mathbf{v}, \tag{89}$$

where the external force is composed of a driving by myosin motors, $\mathbf{F}_{myosin}$ and a frictional drag, $\gamma\mathbf{v}$, leading to a fluid velocity $\mathbf{v}$. Note that the units of all of these terms are $force/volume$. Next, we compare the relative magnitudes of the friction and viscous terms to determine if our system is indeed in the "dry" active matter regime in which

$$\gamma\mathbf{v} \gg \eta\nabla^2\mathbf{v}. \tag{90}$$

If so, friction is quantitatively much stronger than the viscous drag due to interactions between adjacent fluid elements. We can rewrite that condition as

$$\frac{\gamma}{\eta} \gg \frac{\nabla^2\mathbf{v}}{\mathbf{v}}. \tag{91}$$

To understand the source of friction in our two-dimensional system and estimate its magnitude, we must remember that our two-dimensional flow is in fact an approximation of a very thin, but three-dimensional flow as shown in Figure S8. Instead of a thin 3D fluid layer with viscous internal forces in all three directions, we have a two-dimensional flow in which internal viscous interactions across the third

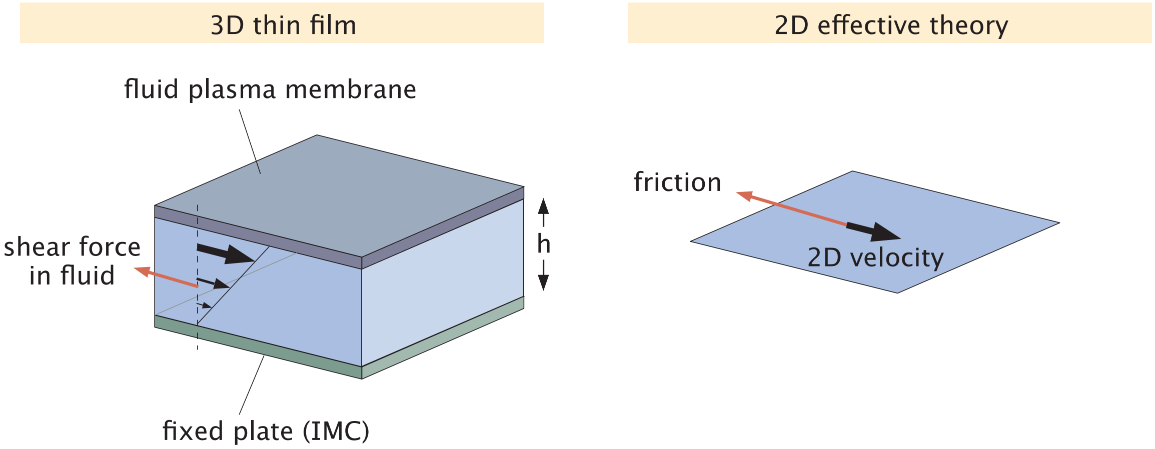

**Figure S8.** Effective theory of two-dimensional fluid motion. In the *Toxoplasma* cell, the actin filaments are moving in a thin layer between the plasma membrane and the inner membrane complex. That thin layer of fluid motion results in internal shear stresses. The effective theory used in our Toner-Tu description is an idealized two-dimensional fluid where the fluid coupling to the no slip boundary conditions in the 3D thin film is replaced by a friction force. Essentially, viscous stresses in the direction perpendicular to the inner membrane complex, idealized here as a fixed plate, are replaced with an in-plane friction.

dimension, $x_3$, are captured as a friction with the fixed substrate. To estimate this friction coefficient, we approximate the viscous term in $x_3$ using

$$\frac{\text{force}}{\text{volume}} = \frac{\partial \sigma}{\partial x_3} \approx \frac{\sigma}{h} \approx \eta \frac{v}{h^2}. \tag{92}$$

Equating this internal shear force in $x_3$ with a two-dimensional frictional drag allows us to make the approximation

$$\eta \frac{v}{h^2} = \gamma v \implies \gamma = \frac{\eta}{h^2} \approx \frac{10^{-3}\,\mathrm{N \cdot s/m^2}}{(10^{-8}\ \mathrm{m})^2} \approx 10^{13} \frac{\mathrm{N \cdot s}}{\mathrm{m^4}}, \tag{93}$$

providing an estimate for the friction coefficient in terms of the known viscosity and the thickness of the fluid layer.

We can now check to see whether we are in the regime of dry active matter implied by eqn. 90. Using $10^{-3}\,\frac{\mathrm{N \cdot s}}{\mathrm{m^2}}$, the viscosity of water, as a reasonable estimate for $\eta$, and plugging in the relevant numbers, we find the scale of frictional drag to be

$$\gamma \mathbf{v} \approx \left(10^{13} \frac{\mathrm{N \cdot s}}{\mathrm{m^4}}\right) \left(10^{-6} \frac{\mathrm{m}}{\mathrm{s}}\right) \approx 10^{7} \frac{\mathrm{N}}{\mathrm{m^3}} \tag{94}$$

and the scale of viscous forces to be

$$\eta \nabla^2 \mathbf{v} \approx \left(10^{-3} \frac{\mathrm{N \cdot s}}{\mathrm{m^2}}\right) \left(\frac{10^{-6}\ \mathrm{m/s}}{(10^{-6}\ \mathrm{m})^2}\right) \approx 10^{-3} \frac{\mathrm{N}}{\mathrm{m^3}}. \tag{95}$$

Thus,

$$\gamma \mathbf{v} \gg \eta \nabla^2 \mathbf{v}, \tag{96}$$

consistent with the dry active matter limit entertained here and the choice of a Toner-Tu model.

*10.2. Estimates of F-actin polymerization and depolymerization rates*

To get a feeling for the polymerization (filament production) rate $c$, we consider actin dynamics in better-studied systems, where we can use published measurements and the BioNumbers database [59] to perform better-informed order-of-magnitude estimates.

First, we consider the comet tail behind motile *Listeria monocytogenes* bacteria as they hijack the host cell actin machinery to propel themselves at a speed that is dictated by their comet tail actin dynamics. For this estimate, we assume a modest speed of 1/10 of a body length each second, or $v \approx 200$ nm/s. We estimate that the propulsion is due to a collection of aligned actin filaments behind the bacterium with a mean spacing of $\approx 50$ nm [60] resulting in a comet tail with a cross-sectional profile of $20 \times 20 = 400$ filaments, or a 1000 nm $\times$ 1000 nm area. We assume that each of these filaments is 100 nm in length. Thus, to move at a speed of $\approx 200$ nm/s (2 filaments/s), the total filament polymerization rate must be

$$\text{actin production rate} \approx 2 \times 400 \, \frac{\text{filaments}}{\mu m^2 \cdot s}. \tag{97}$$

Using this result, we make the order-of-magnitude estimate for the *Toxoplasma gondii* actin polymerization rate of

$$c \approx 10^3 \frac{\text{filaments}}{\mu m^2 \cdot s}, \tag{98}$$

implying a steady-state density

$$\rho_{ss} = \frac{c}{\gamma} \frac{A_{conoid}}{A_{tot}} \approx \frac{10^3 \, \frac{\text{filaments}}{\mu m^2 \ s}}{0.5 \ s^{-1}} \frac{1}{20} \approx 100 \, \frac{\text{filaments}}{\mu m^2}. \tag{99}$$

This number matches our estimated critical filament density, $\rho_c \approx 100 \ \mu m^{-2}$.

Considering an actin comet tail of steady-state length $\approx 5$ $\mu$m, we can also estimate a filament lifetime of $1/\gamma \approx 5 \ \mu m \ / \ 0.2 \ \mu m \ s^{-1} \approx 25$ s. Since actin filaments in *Listeria* comet tails are stabilized and bundled by actin-binding proteins [61], this estimate of $\gamma \approx 0.04 \ s^{-1}$ likely represents a far lower bound on filament turnover rate for *Toxoplasma gondii*, where filaments are not thought to form stable bundles, and the activity of actin depolymerizing factor (ADF) is important to gliding motility [10].

For another simple estimate of depolymerization rate, we can consider actin off rates, measured *in vitro* to be $\approx 7$ monomers/s [62]. Given a filament length of 100 nm $\approx 14$ monomers, we estimate a filament lifetime for *Toxoplasma* actin of

$$\text{filament lifetime} = \frac{1}{\gamma} \approx \frac{14 \text{ monomers}}{7 \text{ monomers/s}} \approx 2 \ s, \tag{100}$$

or a filament turnover rate of $\gamma \approx 0.5 \ s^{-1}$, which matches our estimates for $\gamma$ in Section 7.2. We expect proteins such as ADF to increase filament turnover in *Toxoplasma gondii*, further increasing $\gamma$. Thus, our final estimate for a reasonable range for *Toxoplasma gondii* filament lifetime is between 0.2 s and 10 s, which is equivalent to a range of filament turnover rate $\gamma$ between 0.1 filaments/s and 5 filaments/s. We note that cells may actively tune depolymerization rate over an order of magnitude through the activity of proteins like ADF [63].
